# Mapping Somatosensory Afferent Circuitry to Bone Identifies Neurotrophic Signals Required for Fracture Healing

**DOI:** 10.1101/2024.06.06.597786

**Authors:** Mingxin Xu, Neelima Thottappillil, Masnsen Cherief, Zhao Li, Manyu Zhu, Xin Xing, Mario Gomez-Salazar, Juliet M. Mwirigi, Ishwarya Sankaranarayanan, Diana Tavares-Ferreira, Chi Zhang, Xue-Wei Wang, Mary Archer, Yun Guan, Robert J. Tower, Patrick Cahan, Theodore J. Price, Thomas L. Clemens, Aaron W. James

## Abstract

The profound pain accompanying bone fracture is mediated by somatosensory neurons, which also appear to be required to initiate bone regeneration following fracture. Surprisingly, the precise neuroanatomical circuitry mediating skeletal nociception and regeneration remains incompletely understood. Here, we characterized somatosensory dorsal root ganglia (DRG) afferent neurons innervating murine long bones before and after experimental long bone fracture in mice. Retrograde labeling of DRG neurons by an adeno-associated virus with peripheral nerve tropism showed AAV-tdT signal. Single cell transcriptomic profiling of 6,648 DRG neurons showed highest labeling across CGRP+ neuron clusters (6.9-17.2%) belonging to unmyelinated C fibers, thinly myelinated Aδ fibers and Aβ-Field LTMR (9.2%). Gene expression profiles of retrograde labeled DRG neurons over multiple timepoints following experimental stress fracture revealed dynamic changes in gene expression corresponding to the acute inflammatory (*S100a8*, *S100a9*) and mechanical force (*Piezo2*). Reparative phase after fracture included morphogens such as *Tgfb1, Fgf9* and *Fgf18*. Two methods to surgically or genetically denervate fractured bones were used in combination with scRNA-seq to implicate defective mesenchymal cell proliferation and osteodifferentiation as underlying the poor bone repair capacity in the presence of attenuated innervation. Finally, multi-tissue scRNA-seq and interactome analyses implicated neuron-derived FGF9 as a potent regulator of fracture repair, a finding compatible with in vitro assessments of neuron-to-skeletal mesenchyme interactions.

## Main

Primary somatosensory neurons comprise a diverse subset of neurons, which communicate information about the external environment and internal state to the central nervous system enabling perception and reaction to a wide range of stimuli including pain^1^. These somatosensory neurons have likewise been shown to be essential for processes of tissue repair and regeneration across species and organ systems^2^. In clinical contexts, peripheral neuropathy, caused either by trauma, cancer treatment or disease, is associated with tissue atrophy and poor wound healing^3^.

In tissue repair, the somatosensory neurons regulates diverse cell processes such as cell recruitment and migration^4^, cell proliferation, differentiation^5,6^, and immune responses^7^, principally via paracrine factor release.

During development, the skeleton is richly innervated by the Tropomyosin receptor kinase A (TrkA) expressing sensory neurons which accounts for the profound pain following bone fracture^8,9^. Most prior work on skeletal neurobiology has focused on understanding bone nociceptive pathways, but recent studies provide evidence that sensory nerves also function to initiate bone formation during skeletal morphogenesis. For example, TrkA sensory nerves innervate the developing skeleton in close temporal association with and proximity to sites of incipient bone formation^8,10^. Such data suggest the possibility that somatosensory nerves release bone inductive signals required for skeletal stem cell proliferation and differentiation. Strikingly, these experimental observations have recently been reflected in clinical trials for the FDA-approved Trk inhibitor Entrectinib, which is associated with an increased risk of bone fracture^11^. Whether or to what extent such bone morphogenic and nociceptive actions are mediated by distinct neuronal pathways has been difficult to study in part due to lack of tractable model systems for studying nerve-bone interactions and the extensive heterogeneity of peripheral sensory neurons^12–14^.

In this study we validated a mouse model for visualizing and characterizing signaling of sensory nerves that innervate the skeleton. Using this approach, we mapped the anatomical features of the DRG-bone sensory nerve circuitry and characterized the temporal transcriptomic landscape of DRG neurons innervating long bone fractures at single cell resolution. In this manner, we have uncovered neural-skeletal interactions that underlie neuron-guided tissue repair.

## Results

### Retrograde labeling identifies unique skeletal-innervating sensory neurons

In order to identify peripheral sensory neurons that innervate bone, an engineered adeno-associated virus with peripheral nerve tropism (AAV-PHP.S-tdTomato, AAV-tdT)^15^ was used to retrograde label dorsal root ganglia (DRG) neurons. AAV-tdT signal was examined every week from 1wk until 4 wks after periosteal injection at midshaft ulna. AAV-tdT signal was detected in the expected ipsilateral C7-T1 DRGs innervating the forelimb from 1 wk and progressively increased over 4 wks, also confirmed with fast blue labeling (**Fig. 1a-b and Extended Data Fig. 1**a), while little or no labeling was observed within distant ipsilateral L1 DRGs (**Extended Data Fig. 1**b). In order to characterize the transcriptomic profile of skeletal-innervating sensory neurons, DRGs were harvested and subjected to scRNA-sequencing (scRNA-seq) after labeling. Sequencing recovered 1,020 neurons. Neurons were classified into 14 different clusters using previously established molecular taxonomy^12^ and according to gene marker expression (**Fig. 1c, d**). Viral labeling was determined by expression of AAV-tdT reporter gene *AAV-gene1*. AAV labeling of skeletal-innervating neurons by subcluster was next evaluated, with the highest labeling found across CGRP^+^ neuron clusters (5.7-11.1% across individual clusters) and Aβ- Field LTMR (5.4%) (**Fig. 1e**). These neurons belong to unmyelinated C fibers (CGRP-β/γ, CGRP-ε: Tac1-expressing peptidergic nociceptors), thinly myelinated Aδ fibers (CGRP-ζ,

**Fig. 1.**
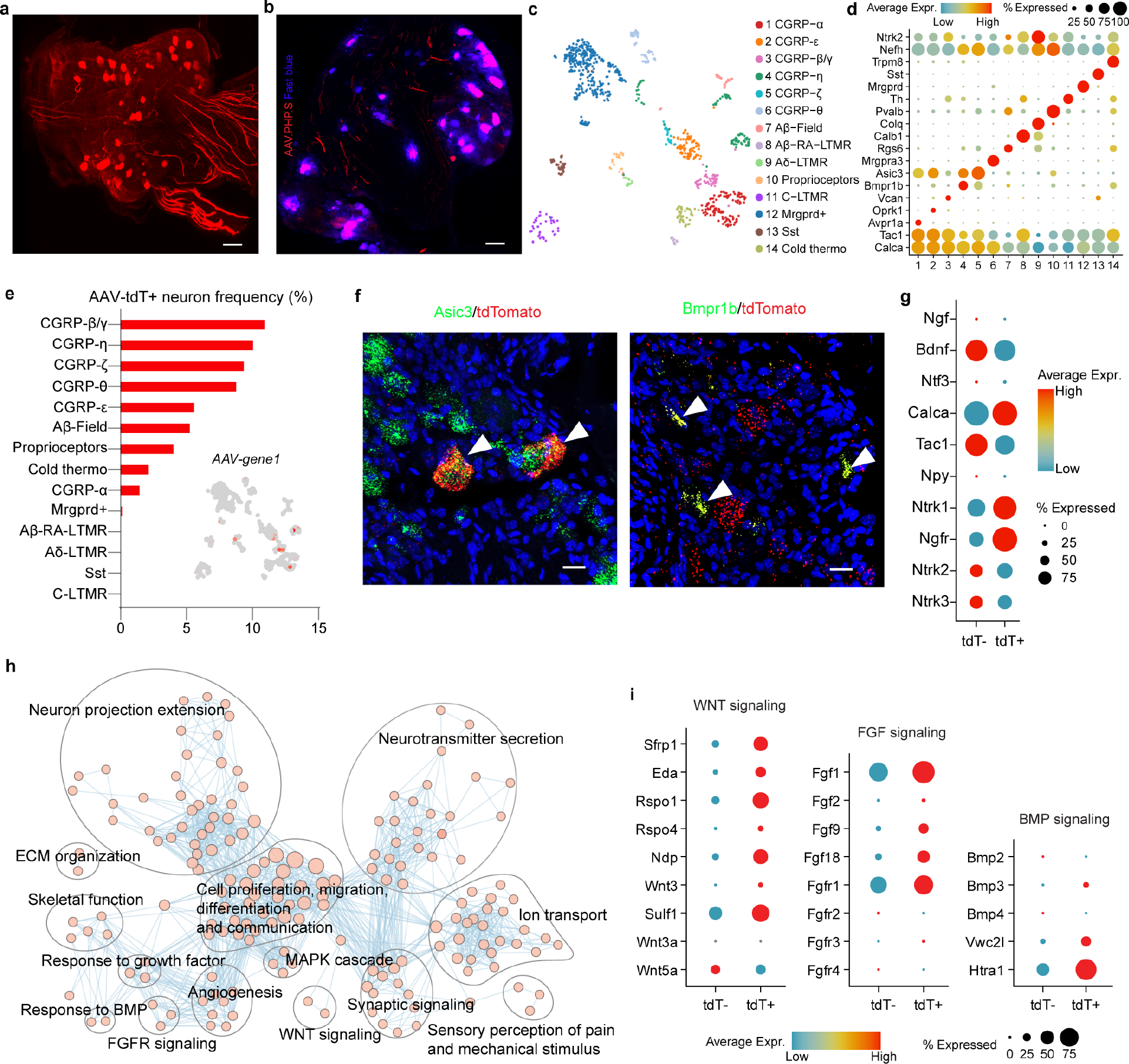
Identification and characterization of skeletal-innervating sensory neurons. Periosteal injection of AAV-PHP.S was performed into the ulnar mid-diaphysis of 14 wk old C57BL/6J male mice, following by analysis of ipsilateral innervating dorsal root ganglia (DRG) after 4 wks labeling. **a** Whole mount imaging of tdTomato labeling of C7 DRG by IHC. **b** Co-visualization of tdTomato and fast blue labeling in ipsilateral C7 DRG. Inset shows contralateral DRG. **c** UMAP of all neuron clusters (N=1,020 neurons). **d** Dot plot of marker gene expression for each neuron cluster. **e** Frequencies of AAV-tdT labeling by neuron cluster, inset shows UMAP of neuronal AAV labeling identified by *AAV-gene1*. **f** RNAscope co-localization of tdTomato with either *Asic3* or *Bmpr1b*, markers for CGRP-ζ and CGRP-η neurons, respectively. **g** Dot plot of neurotrophins, neurotrophin receptors and neuropeptides among tdT^+^ and tdT^-^ neurons. **h** Enrichment map of significantly enriched gene ontology (GO) in tdT^+^ neurons as compared to tdT^-^ neurons. Nodes represent gene-sets and edges represent GO defined relations. Clusters are annotated according to the corresponding function. Enrichment map with complete labeling is shown as Extended Data Fig. 1c. **i** Dot plots of signaling components significantly expressed in tdT^+^ neurons as compared to tdT^-^ neurons for WNT, FGF and BMP signaling. Data obtained from N=12 DRGs (C7, C8 and T1) from N=4 mice. Scale bar: 100μm for **a** and **b**, 20μm for **g**.

CGRP-η: Ntrk1^+^ and Nefh^+^) and Aβ-Field LTMR^1,16,17^. Negligible AAV labeling was observed in clusters known for primarily cutaneous innervation, such as *Trpm8*^+^ cold thermoceptors, *Mrgprd*^+^ nonpeptidergic nociceptors, and *Sst*^+^ pruriceptors. *In situ* hybridization confirmed expression of tdT labeling in representative CGRP^+^ clusters, including *Asic3*^+^ CGRP-ζ neurons and *Bmpr1b*^+^ CGRP-η neurons (**Fig. 1f**).

Characterization of AAV-tdT labeled skeletal-innervating neurons was next performed. In keeping with our and others past reports^9,18,19^, tdT^+^ neurons showed high levels of the high affinity NGF receptor *Ntrk1,* but also the low-affinity receptor *Ngfr* (**Fig. 1g**). In comparison, higher expression of *Ntrk2* and *Ntrk3* was found on tdT^-^ neurons. Neuropeptides and neurotrophins displayed differential expression among labeled versus unlabeled neurons. *Calca* was highly expressed in tdT^+^ neurons, whereas *Bdnf, Npy and Tac1* were more highly expressed in tdT^-^ neurons (**Fig. 1g**). Characterization of labeled neurons was performed by Gene Ontology (GO) enrichment analysis (**Fig. 1h, Extended Data Fig. 1**c). In comparison to unlabeled neurons, tdT^+^ neurons showed enrichment for terms related to the regulation of skeletal functions (e.g. osteoblast and chondrocyte differentiation, skeletal development, angiogenesis) as well as a host of intrinsic neuronal functions (e.g. neuron projection morphogenesis, neurotransmitter secretion, synaptic signaling). In addition, tdT^+^ neurons showed enrichment for several signaling pathways, including fibroblast growth factor (FGF), Wnt, bone morphogenetic protein (BMP) and mitogen-activated protein kinase (MAPK) signaling (**Fig. 1i, Extended Data Fig. 1**c). Secreted nerve factors related to enriched signaling pathways were also identified, such as genes related to FGFR signaling (*Fgf1*, *Fgf2*, *Fgf9*, *Fgf18*), Wnt signaling (*Sfrp1*, *Eda*, *Wnt3*, *Ndp*, *Rspo1*, *Rspo4*) and BMP signaling (*Bmp3*) (**Fig. 1i**) (See **Supplementary Information File 1** for a full list of secreted genes significantly upregulated in tdT^+^ neurons). To exclude the possibility that the observations above were due to relative enrichment for CGRP^+^ neuron clusters, a focused comparison between labeled and unlabeled neurons within the three top-labeled CGRP- expressing neuron clusters was performed, with similar GO enrichments (**Extended Data Fig. 1**d).

### Fracture induces a dynamically regulated temporal response within sensory neurons

To demonstrate the temporal responses of sensory neurons to bone tissue injury, scRNA-seq was obtained from retrograde-labeled DRG tissues at 3 different timepoints after stress fracture using a previously validated ulnar stress fracture model^19^ and compared to the uninjured condition (**Fig. 2a**). Timepoints of d1, d14 and d56 post-fracture were chosen to capture the acute inflammatory, reparative and remodeling phases of bone tissue repair, respectively. A similar composition of neuron cell clusters was seen among injured timepoints in comparison to the uninjured timepoint (**Extended Data Fig. 2**). A slight change of order was seen in frequencies of neuron labeling per cluster, with Aβ-RA-LTMR neurons more represented (**Extended Data Fig. 2**h). A total of 389 tdT^+^ neurons and 16,583 genes were evaluated for temporal patterns after injury.

**Fig. 2.**
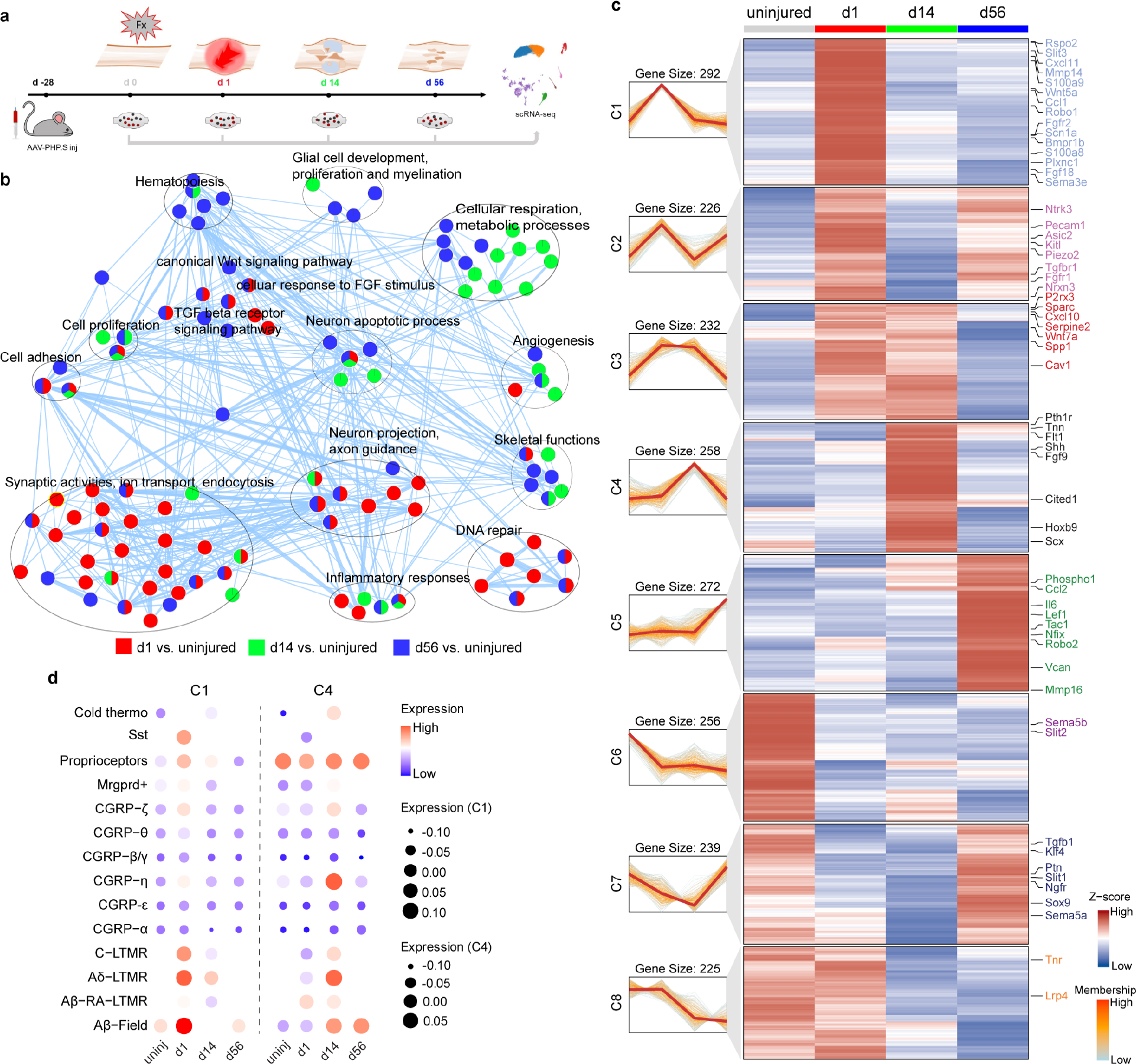
Temporal responses of sensory neurons to fracture injury. a. Schematic diagram of the experiment, in which mice were subjected to periosteal labeling by AAV.PHP.S 4 wks prior to stress fracture. Stress fracture was performed in 18 wks old mice. Ipsilateral C7-T1 DRGs were harvested for scRNA-seq within uninjured mice, as well as 1, 14, and 56 d post-fracture. N=12 DRGs from 4 mice for uninjured, 18 DRGs for 6 mice per injured timepoint. **b** Enrichment map displays the significantly enriched gene-sets in tdT^+^ neurons after fracture vs. uninjured condition. Nodes represent gene-sets and edges represent GO defined relations. Clusters are annotated according to the corresponding function (red represents GO terms significantly enriched in d1, green represents GO terms significantly enriched in d14, blue represents GO terms significantly enriched in d56). Enrichment map with complete labeling is shown as Extended Data Fig. 3a. **c** Soft clustering of 2,000 highly variable genes across four different timepoints by Mfuzz using ClusterGvis in tdT^+^ DRG neurons. N= 389 neurons in total. **d** Dot plots of module scores comprised of genes from cluster C1 and C4 across time points stratified by cell types in tdT+ neurons.

Pseudobulk sequencing analysis was used to examine biological processes in the neuronal response to fracture repair and dynamic changes across time (**Fig. 2b, Extended Data Fig. 3**a**, Supplemental Information File 2**). Skeletal-innervating sensory neurons 1 d after fracture showed enrichment for GO terms related to inflammatory responses, synaptic activities, ion transport and endocytosis. At 14 d after injury, GO term enrichment included groups of terms related to tissue regeneration including cell proliferation, angiogenesis, skeletal functions as well as metabolic processes. At d56 after fracture, upregulated GO terms were associated with hematopoiesis and glial cell development and proliferation.

Eight different gene sets were identified in tdT^+^ neurons by Mfuzz according to their dynamic expression (**Fig. 2c**, **Supplemental Information File 3**). Acute responders (C1-C2, those increase at d 1 after fracture) include genes which play a role in sensory perception of pain (*Scn1a* and *Kcna1*) and mechanical detection (*Piezo2*, *Asic2*). C1 genes also include regulators of inflammation (*S100a8*, *S100a9*, *Cxcl11*) and axon guidance molecules and receptors (*Sema3e*, *Slit3*, *Wnt5a*, *Robo1*) which peaked on d1 and gradually went down to baseline. C3 and C4 clusters include genes that upregulated on d14, including genes regulating skeletal system development, osteoblast/chondrocyte differentiation and ECM regulation (*Shh*, *Fgf9*, *Pth1r*, *Scx*, *Wnt7a*, *Spp1*, *Sparc*), as well as angiogenesis (*Flt1*, *Cxcl12*, *Cav1*). C5 genes are a group of genes that peaked at d56, including genes related to bone mineralization such as *Phospho1* and *Nfix*, as well as genes related to resolution of inflammation and would healing, such as *Sema7a* and *Tnc*. In addition, C6, C7 and C8 represent genes that largely fell in expression after injury.

For example, repellant cues for axon growth such as *Sema5a*, *Sema5b*, *Slit1*, *Slit2* and *Tnr* were decreased. *Ptn* levels were relatively higher at uninjured and d56 and lower at d1 and d14, consistent with previous study showing increased PTN in bone lead to impaired fracture healing^20^. A full list of gene sets is shown in **Supplemental Information File 3**. To identify which labelled neuron clusters were primarily responsible for the transcriptomic changes post-fracture, module scores comprised of C1 and C4 gene sets were analyzed (corresponding to expression peaks at d1 and d14, respectively). Dot plot revealed that C1 gene set expression changes were seen across multiple neuron clusters, but primarily in Aβ-Field, Proprioceptors, CGRP-ζ and CGRP-η cells (**Fig. 2d**). In a similar manner, C4 gene set expression changes were primarily observed in Aβ-Field, CGRP-ζ and CGRP-η cells (**Fig. 2d**).

For comparison, unlabeled (tdT^-^ neurons) were next evaluated in a similar manner (**Extended Data Fig. 3**b, **Supplemental Information File 4**). In tdT^-^ neurons, when compared to the uninjured condition, enrichment in terms related to immune response were seen in all injured time points. At 14 d after injury, some similarity in upregulated GO terms was observed (for example similar induction of terms related to angiogenesis and cell proliferation were seen), however no terms related to skeletal function-related terms were seen. Distinct gene patterns were also identified in tdT^-^ neurons (**Extended Data Fig. 3**c), suggesting distinct responses to fracture injury between skeletal-innervating neurons as compared to all other somatosensory neurons within the same dermatomyosclerotome.

Finally, gene regulatory network analysis also uncovered unique patterns of transcription factor (TF) activity changes across each phase of tissue repair in tdT^+^ neurons (**Extended Data Fig. 3**d). At d1 post-fracture, *Zfp263*, *Nfkb2* and *Stat1* were significantly upregulated, TFs associated with the regulation of inflammation in other systems (*Nfkb2* and *Stat1*^21–23^). At d14 post-fracture, several TFs were identified, including regulators of neural development (*Maz* and *Gata2)*^24,25^, NGF target genes (*Egr3*)^26,27^, negative regulators of neuronal apoptosis (*Klf2*)^28^, as well as stress-responsive genes (*Atf4*)^29^. Interestingly, common neuronal markers of nerve injury (e.g. such as *Atf3, Jun, Klf6, Pou2f1* and *Sox11*) were not conspicuously increased after bone fracture, in comparison to established nerve injury models^13^ (**Extended Data Fig. 3**e).

### Denervation impairs fracture healing via alterations in mesenchymal cell proliferation and differentiation

To evaluate how denervation affects fracture healing at single cell resolution, two denervation models were utilized in parallel. First, a previously validated chemical-genetic approach was used, in which knockin of a point mutation in TrkA allows for inhibition of TrkA catalytic activity via the small molecule 1NMPP1 (TrkA^F592A^ mice)^19,30^. Next a surgical neurectomy procedure was performed, in which the ulnar nerve was transected above the level of the Olecranon process prior to fracture. Validation of surgical neurectomy was performed, in which both intra-epidermal and ulnar periosteal nerves showed a significant reduction in frequency (64- 88% across sites and neural antigens, **Extended Data Fig. 4**a-d). In addition, neurectomy led to a severe diminishment in fracture callus size, including a 57.2% reduction in bone volume (BV) and 62.4% reduction in tissue volume (TV) (**Fig. 3a**), findings similar to those we previously reported in TrkA^F592A^ mice^19^. Histologic analysis showed a smaller callus area by H&E, confirmed by less cartilage and bone by Safranin O/Fast Green (SO/FG) and alkaline phosphatase (ALP), respectively (**Extended Data Fig. 4**e). Next, to understand how denervation affects the transcriptomic profiles of callus resident cells, scRNA-seq was performed on callus tissues 14 d after fracture. Two parallel experiments were performed (**Fig. 3b**), in which fracture healing was monitored at the single cell level in the context of either surgical or chemical-genetic denervation. First, ulnar neurectomy was performed in comparison to sham surgery alone.

**Fig. 3.**
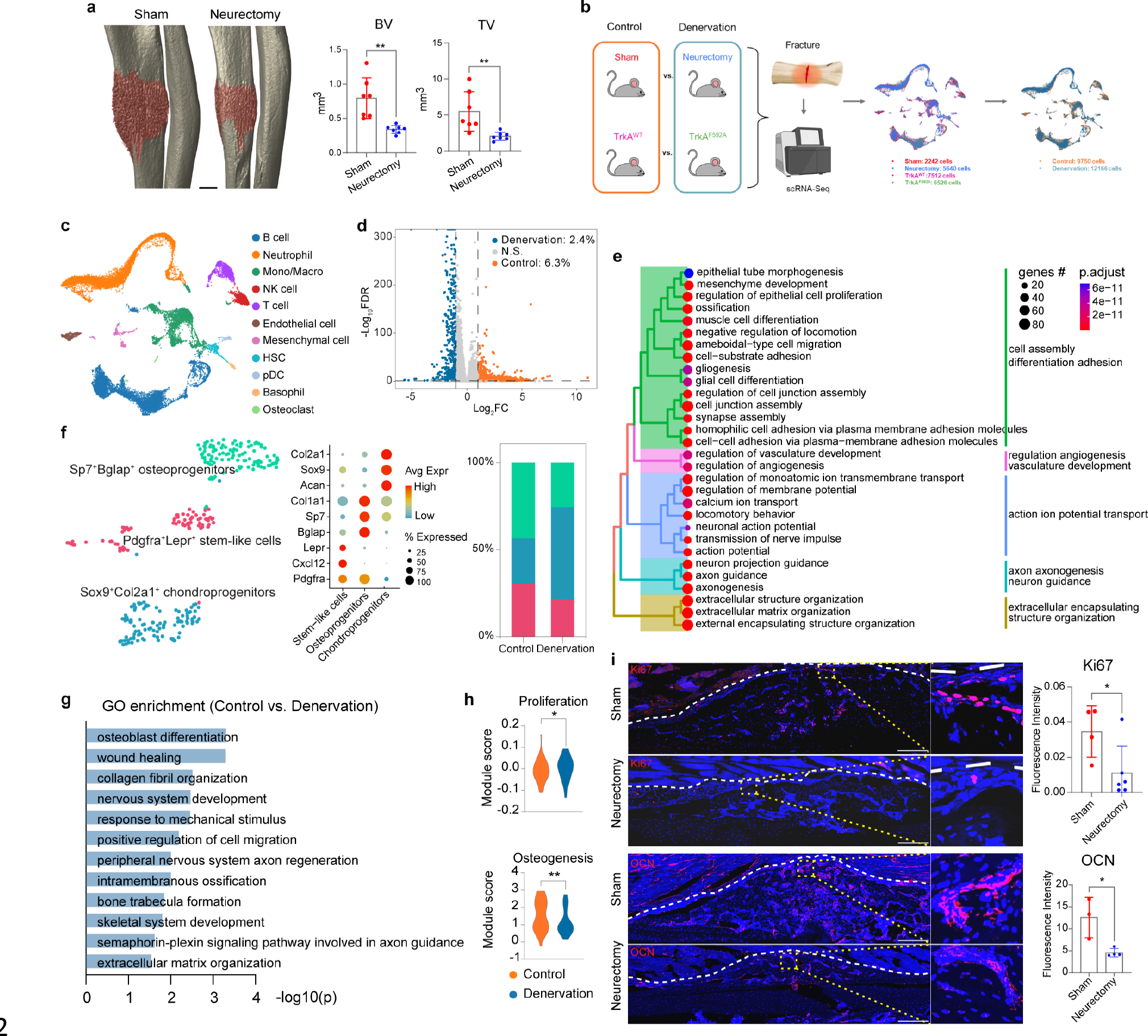
Denervation impairs fracture repair at gross and single cell resolution. a. Left: Representative μCT images of fracture callus from sham-operated and neurectomized animals. Fracture callus colorized brown. Right: Quantification of fracture callus Bone Volume (BV) and Tissue Volume (TV). **b** Experimental schematics of two parallel experiments via either surgical or chemical-genetic denervation. Surgical denervation was achieved via ulnar nerve transection, sham surgery was performed as a control. Chemical-genetic denervation was achieved by 1NMPP1-treated TrkA^F592A^ mice, 1NMPP1-treated TrkA^WT^ mice as a control. Stress fracture was performed on mice from both experiments at 18 wks old and callus tissue was harvested at d14 post-fracture for scRNA-seq. **c** UMAP of callus cells from all groups by cell cluster. **d** Volcano plot of overrepresented genes in control and denervation groups. Significantly differentiated expressed genes are identified using thresholds with FDR < 0.05 and Log2FC > 1. Separate comparison for each experiment shown in Extended Data Fig. 6-7. **e** Tree plot of GO enrichment in control groups in comparison to denervation groups. Additional separate comparisons shown in Extended Data Fig. 6-7. **f** Left: UMAP of mesenchymal cell subclusters. Middle: Dot plot of marker gene expression in each mesenchymal subcluster. Right: Cell frequency of mesenchymal cell subclusters in control and denervation groups. **g** GO enrichment in mesenchymal cells in control groups in comparison to denervation groups. **h** Module score analysis of proliferation (among cells of early pseudotime: 0-1), and osteogenesis (among cells of later pseudotime: >2) between control groups and denervated groups of mesenchymal cells. **i** Immunofluorescent staining and quantification of Ki67 and Osteocalcin (OCN) in sham-operated and neurectomized fracture calluses. N=3-6 mice per group. *p<0.05; **p<0.01. Scale bar in **a**: 500μm, **j**: 200μm. Values plotted are the means with errors bars representing ± 1 s.d.

Second, 1NMPP1-treated TrkA^F592A^ mice were examined in comparison to 1NMPP1-treated TrkA^WT^ mice. Sequencing of fracture callus tissue retrieved in total 21,920 cells, with 9,754 cells in control groups (Sham: 2,242 cells, TrkA^WT^: 7,512 cells) and 12,166 cells in denervation groups (Neurectomy: 5,640 cells, TrkA^F592A^: 6,526 cells) (**Extended Data Fig. 5**a). Cells were then grouped into 11 clusters according to the expression of marker genes (**Fig. 3c, Extended Data Fig. 5**b). Cells derived from the denervation group were associated with a lower frequency of mesenchymal cells and endothelial cells, and higher frequency of neutrophils; however all cell clusters were represented across all conditions (**Extended Data Fig. 5**c). Using pseudobulk sequencing, of a total of 20,941 genes, 1,327 genes (6.3%) were overrepresented under control conditions, while 503 genes (2.4%) were overrepresented under denervation conditions (**Fig. 3d**, **Extended Data Fig. 6**a**, Extended Data Fig. 7**a **for separate comparison, Supplemental Information File 5**). GO term enrichment using all callus resident cells was next performed.

Within the control groups, enrichment for terms related to cell differentiation and adhesion, angiogenesis, ion transport, axonogenesis and extracellular structure organization were observed (**Fig. 3e**). Similar results were seen when comparing the two denervation models separately (**Extended Data Fig. 6**b**, Extended Data Fig. 7**b). GO terms enriched in the denervation groups were mostly related to immune responses, observed even after correction for cluster frequency (**Supplemental Information File 6**).

A focused examination of mesenchymal cells within the fracture callus was next performed (**Fig. 3f-i**). A total of 295 mesenchymal cells were further classified into three subclusters based on characteristic gene markers into *Pdgfra*^+^*Lepr*^+^ stem-like cells, *Sp7*^+^*Bglap*^+^ osteoprogenitors and *Sox9*^+^*Col2a1*^+^ chondroprogenitors (**Fig. 3f**). Among all mesenchymal cells, GO terms enriched in the control groups were related to bone formation such as osteoblast differentiation, wound healing, collagen fibril organization, intramembranous ossification and bone trabecula formation when compared to the denervation group (**Fig. 3g**) (**Supplemental Information File 7-9**).

Pseudotemporal analysis revealed a disturbed gene expression pattern in mesenchymal cells derived from the denervation groups (**Extended Data Fig. 8**a), with denervation groups showing enrichment in genes related to chondrogenesis in late pseudotime (e.g., *Sox9, Comp, Acan* and *Col2a1*) while control groups showing some enrichment in genes associated with stemness in early pseudotime (e.g., *Lepr, Ebf3* and *Cxcl12*) and osteogenesis in late pseudotime (e.g., *Bmp1* and *Col1a1*). To discern the cellular effects of denervation more quantitatively on compromised bone-forming functions in mesenchymal cells, we evaluated a range of discrete mesenchymal cell activities, including stemness, proliferation, apoptosis, migration and differentiation. Module scores of genes related to cell stemness, migration or apoptosis showed no significant differences between mesenchymal cells across experimental conditions (**Extended Data Fig. 8**b). In contrast, module scores of gene sets related to cellular proliferation and osteogenic differentiation showed significant reductions in mesenchymal cells derived from the denervation groups (**Fig. 3h, Supplemental Information File 10**). These changes in proliferation and osteogenesis were further validated using Ki67 and osteocalcin (OCN) immunohistochemistry (IHC) staining, with a 71.1% decrease in Ki67 and 63.7% decrease in OCN immunostaining detected (**Fig. 3i**) in the context of denervation conditions. These results suggest that skeletal-innervating neurons normally play a role in the promotion of fracture repair via the positive regulation of proliferation and osteogenic differentiation of periosteal mesenchymal cells *in vivo*.

### Neural derived FGF9 induces skeletal repair via modulation of periosteal cell proliferation and differentiation

To further decipher the neural-skeletal interactions that may regulate fracture repair, interactome analysis was performed to infer intercellular communications. CellChat analysis predicted that DRG neurons, as a signaling source, interact with every callus cell type, with the highest number of interactions with mesenchymal and endothelial cells (**Fig. 4a**). Focusing specifically on DRG neuron to mesenchymal cell signaling interactions, 21 signaling pathways interactions were predicted, with PTN, FGF and SEMA3 pathways showing the highest probability of interaction (**Fig. 4b**). Relative information flow was compared between our previously identified tdT^+^ skeletal-innervating neurons vs tdT^-^ neurons (**Fig. 4c**). Results showed that tdT^+^ skeletal- innervating neurons showed enrichment for 13 pathways such as GRN, PERIOSTIN, TGFb, and FGF signaling, among others (**Fig. 4c**). Of these 13 pathways, only 3 pathways were computationally identified to have DRG neurons as the primary sender cell population, including FGF, GAS, and Hedgehog (HH) signaling (**Fig. 4d, Extended Data Fig. 9**a). Other signaling pathways showed callus resident cells as the predominant sender population and were not further pursued as neural-skeletal interactors (**Extended Data Fig. 9**b-i). Nichenet analysis revealed a range of potential ligands and their targets from tdT^+^ neurons that may act on mesenchymal cells at d14 (**Fig. 4e**). The expression of predicted tdT^+^ neural derived ligands were next evaluated in comparison to unlabeled tdT^-^ neurons or callus resident mesenchymal cells, which further narrowed our predicted interactors (**Fig. 4f**). Synthesizing interactome analyses, four individual ligands were identified from FGF and HH signaling pathways as likely neural-derived paracrine regulators, including *Fgf5, Fgf9, Fgf14* and *Shh* (**Fig. 4f**). Of these, only *Fgf9* showed preferential expression in tdT^+^ skeletal-innervating neurons and dynamic temporal transcriptional regulation after fracture repair (**Fig. 4f, g**).

**Fig. 4.**
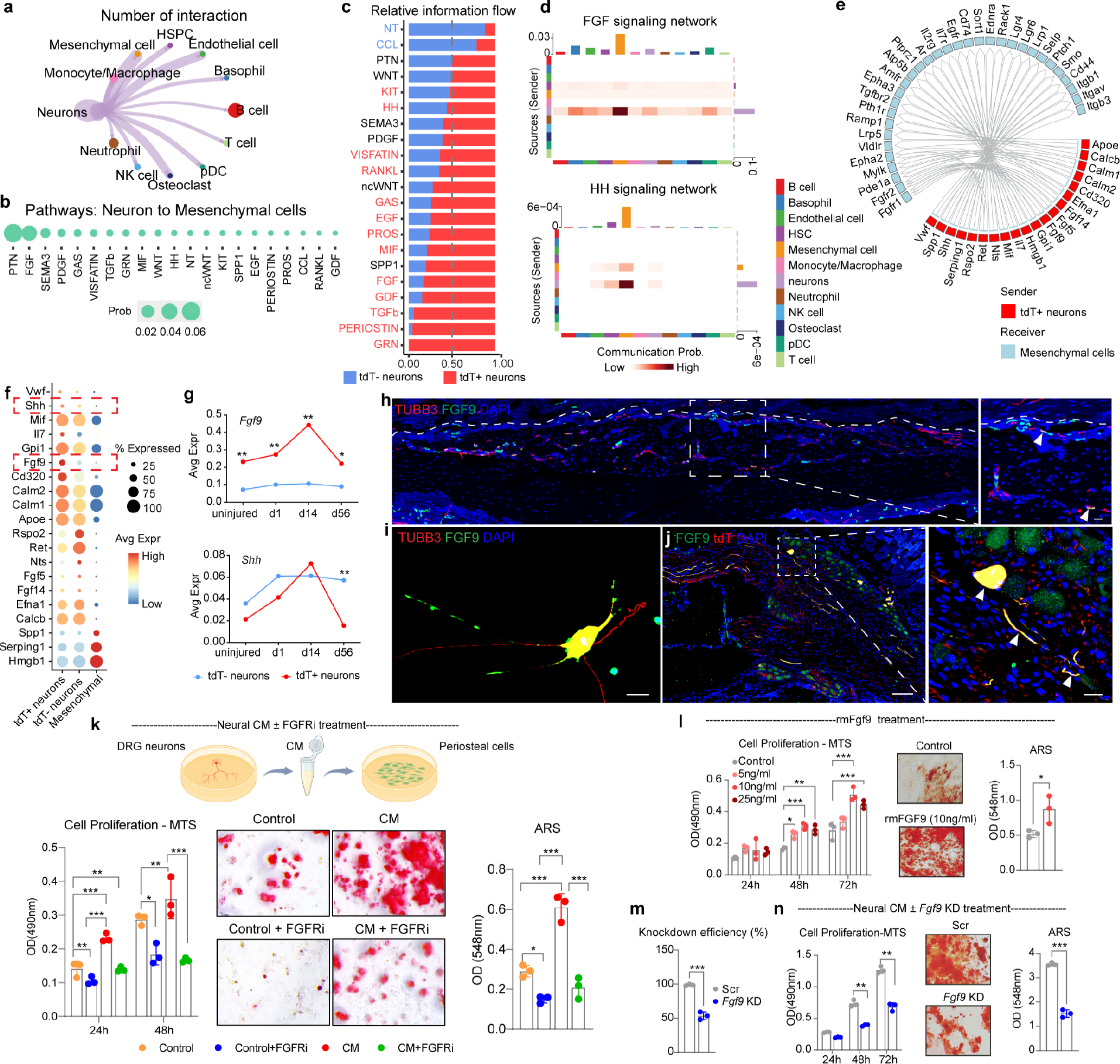
Interaction analyses identify FGF9 signaling in DRG neuronal regulation of fracture callus cell osteogenesis. a. Cell-cell communication network shows the number of interactions between DRG neurons to each callus cell type identified by CellChat (all cells derived from d14 post-injury and control conditions). **b** Dot plot of probability of inferred pathways from neurons to mesenchymal cells (d14 post-fracture cells used). **c** Relative information flow of pathways identified from **b**, from skeletal-innervating neurons (tdT^+^) and non-skeletal- innervating neurons (tdT^-^) to mesenchymal cells. **d** Heatmaps of the number of interactions within FGF, and HH signaling. Only signaling pathways with significant predicted neuronal senders are shown. Other pathways shown in Extended Data Fig. 9. **e** Circos plot visualization to show active ligand-target links from tdT^+^ neurons to mesenchymal cells at d14. **f** Average expression of predicted ligands from **e** by dot plot among tdT^+^ neurons, tdT^-^ neurons and mesenchymal cells. **g** Line graphs of average expression of *Fgf9* and *Shh* within tdT^+^ or tdT^-^ neurons across timepoints. **p<0.01 for comparisons in gene expression between tdT^+^ vs tdT^-^ neurons at each time point. **h** Co-immunostaining for TUBB3 and FGF9 within fracture callus tissue (d14). **i** Co-immunostaining for TUBB3 and FGF9 in cultured DRG neurons. **j** FGF9 and tdT co-immunostaining of T1 DRG after AAV-PHP.S labeling. **k** Top: Experimental schematics of treatment of periosteal cells by DRG neural conditioned medium (CM). Bottom left: MTS assay to assess cell proliferation in periosteal cells treated with control medium, control medium with the FGFR (i)nhibitor PD173074, DRG neural CM and neural CM with PD173074. Bottom middle: ARS staining, 28 d differentiation. Right: Quantification of ARS staining. **l** Left: MTS assay in periosteal cells treated with rmFGF9: 5- 15ng/ml. Middle: ARS staining of control and rmFGF9-treated cells, 28 d differentiation. Right: quantification of ARS to assess osteoblast differentiation of periosteal cells treated with control or 10ng/ml rmFGF9. **m** Knockdown efficiency of *Fgf9* in cultured DRG neurons on d 4 by qRT-PCR from pooled cells of 10 DRGs /per well of 24 well plate. **n** Left: MTS assay in periosteal cells treated with neural CM derived from DRG neurons with Scr or DRG neurons with *Fgf9* siRNA. Middle: ARS staining of periosteal cells treated with neural CM from DRG neurons with Scr or DRG neurons with *Fgf9* siRNA. Right: Quantification of ARS to assess osteoblast differentiation of periosteal cells treated with neuronal CM from DRG neurons with scrambled siRNA (Scr) or DRG neurons with *Fgf9* siRNA. N=3 replicates per group. C57BL/6J mice were used for all the experiments. *p<0.05; **p<0.01; ***p<0.001. Scale bar for left panel in **j**: 100μm, all other scale bars: 20μm. Values plotted are the mean with errors bars representing ± 1 s.d.

Having computationally implicated *Fgf9* as a neural derived paracrine regulator of bone tissue repair, validation was first performed by assessment of FGF9 protein expression. Skeletal-innervating nerves in fracture calluses identified using the pan-neuronal marker TUBB3 showed significant co-localization with FGF9 (**Fig. 4h**). Moreover, FGF9 was detected in paired specimens of human DRGs and sciatic nerves by SOMAscan proteomics assay (**Extended Data Fig. 10**b). Likewise, cultured mouse DRG neurons showed FGF9 immunoreactivity in cell bodies and axons (**Fig. 4i**). In addition, sections of AAV-tdT labeled DRGs confirmed the presence of FGF9 immunoreactivity in tdT^+^ neurons (**Fig. 4j**), and FGFR1-4 likewise were found to be expressed transcriptionally in multiple callus resident cells, including mesenchymal cells (**Extended Data Fig. 10**c). FGF9-FGFR signaling interactions were next formally assessed using mouse DRG neural conditioned medium (CM) applications to mouse periosteal mesenchymal cells in culture (**Fig. 4k**). Neural CM demonstrated a pro-proliferative and pro-osteogenic effect on recipient periosteal cells, which was fully nullified by the co-administration of the pan-FGFR inhibitor (FGFRi) PD173074 (**Fig. 4k, Extended Data Fig. 10**d). Addition of recombinant FGF9 (rmFGF9) to periosteal cell cultures phenocopied neural CM administration, with similar mitogenic and osteogenic effects observed (**Fig. 4l, Extended Data Fig. 10**d).

Finally, siRNA mediated knockdown of *Fgf9* in DRG neurons was performed (**Fig. 4m**) followed by collection of CM. The mitogenic and pro-osteogenic effects of neural CM were significantly diminished among CM derived from *Fgf9* KD neurons (**Fig. 4n, Extended Data Fig. 10**d). In aggregate, our multi-tissue interactome analyses and validation studies confirm the role of FGF9-FGFR signaling in sensory neuron mediated regulation of bone tissue repair capacity.

## Discussion

In this study, we experimentally mapped skeletal-innervating peripheral sensory neurons for the first time with a combined approach of retrograde tracing and scRNA-sequencing. Single cell analyses revealed a temporally dynamic response of DRG neurons to fracture injury, and implicated FGF signaling in neural regulation of bone tissue repair. Composite interactome analyses and validation studies implicated FGF9 from skeletal-innervating neurons as an important regulator of the bone injury repair process.

Bone-innervating neurons as identified in our study primarily encompassed CGRP-expressing C fiber and Aδ fibers, as well as Aβ fibers with low mechanical thresholds. Unsurprisingly, these differ significantly with the neurons that innervate lymph nodes and skin tissue as identified by similar retrograde tracing methods^31^. Bone-innervating neurons express higher levels of NGF receptors *Ntrk1*, *Ngfr* as well as *Calca*, which is in agreement with past reports utilizing immunohistochemistry (IHC) or transgenic reporter mice^8,9^. These nerves are believed to be most sensitive to nociceptive stimuli^32,33^. Previous work by Mantyh *et al* has also shown that mouse femur-innervating nerve fibers express p75 and TrkA and course along nearby NGF^+^ blood vessels^18^. Studies using Prokr2^Advillin^-tdTomato mice also demonstrated that Prokr2-expressing sensory neurons, which corresponds to CGRP-η in our dataset, innervate the tibial periosteum^34,35^. Our results have also shown that skeletal-innervating neurons include Aβ fibers that are responsive to mechanical stimuli, such as Aβ Field-LTMR^1^ and Aβ RA-LTMR (Pacinian corpuscles that innervate the periosteum of bones in rodents^36^). However, whether Pvalb-expressing proprioceptors innervate bone is less clear. In our study, under uninjured conditions, 2.4% of DRG neurons were proprioceptors, of which 4.2% were labeled by the virus.

Proprioceptors are predominately found in muscle and tendon^37^. Studies have shown that the proprioceptive system plays important role in the morphologic restoration of fracture bones^38^, yet no direct evidence to our knowledge has shown the innervation of bone by proprioceptors. Of note, we were not able to co-localize Pvalb^+^ proprioceptors and tdTomato labeling by ISH (data not shown). Thus, the interpretation of Pvalb^+^ proprioceptors directly innervating bone requires further validation.

Fracture invoked a temporal transcriptomic response in DRG sensory neurons, which interestingly recapitulated many features of the bone tissue repair process. At early time points, DRG neurons showed signatures of pain perception and inflammatory responses. At later timepoints, DRG neurons demonstrated transcriptomic changes more characteristic of a regenerative response, including pro-proliferative, angiogenic and osteogenic signals. Multiple past studies have documented the transcriptional response of DRG neurons in neuropathic or inflammatory pain models, including at single cell resolution. In experimental pain models, DRG neurons respond to injury immediately and exhibit prolonged inflammatory responses as late as 21 days post-injury^39,40^. Increased gene expression related to nerve regenerative responses such as cell adhesion molecules and TGF-beta signaling pathway are seen early post-injury^39^. Indeed, axotomy has been characterized to induce mouse neuron type switches and reactive neurons adopt an “injured” *Atf3*^+^ transcriptional state, not seen in inflammatory and chemotherapy- induced pain models^13,16^ nor in our fracture model. Therefore, stress fracture induced a unique response from DRG sensory neurons, associated with a pro-reparative, pro-osteogenic neurosecretory phenotype.

Since the discovery of neural dependent limb regeneration in the salamander two centuries ago^41^, nerve dependence in tissue regeneration and repair has been a phenomenon observed across a wide ranges of species and organ systems. The release of various neurotropic morphogens, growth factors and neuropeptides has been identified as one of the most important mechanisms in nerve dependence, such as Neuregulin-1 (NRG1) in limb and heart regeneration^42,43^, Platelet- Derived Growth Factor (PDGF-AA) and oncostatin M (OSM) in digit tip regeneration^44^, Substance P (SP) and CGRP in skin wound repair^45^, FGF1 and Shh in regeneration of adult mouse incisors^46,47^. Our own prior work also implicated follistatin-like 1 (Fstl1) in the regulation of proliferation and differentiation of bone precursor cells in developing cranium of mice^10^, and vascular endothelial growth factor A (Vegfa) in the regulation of heterotopic ossification^48^. Here and for the first time, we directly profiled skeletal-innervating DRG neurons and identified that FGF9 secreted from sensory neurons promotes the proliferation and osteogenesis of periosteal cells and contributes to bone fracture repair. In our study, denervation *in vivo* and inhibition of FGF9-FGFR signaling *in vitro* were associated with compromised proliferation and osteogenesis in mesenchymal/periosteal cells. However, whether this is due to denervation steering mesenchymal cells into chondrogenesis at the expense of osteogenesis or merely a delay of skeletal cell differentiation warrants further exploration.

Although we showed skeletal-innervating sensory neurons and revealed their role in fracture healing via FGF9-FGFR signaling, our study has limitations. First, stress fracture in our model was induced by mechanical loading to the bone and is a relatively small injury that heals predominantly via intramembranous ossification. This injury type and healing pattern undoubtedly elicits distinct neuronal responses in comparison to other orthopaedic models of bone injury. Second, our study focused on direct neural-periosteal mesenchymal cell interactions in fracture repair. It is entirely possible that neuro-immune^49^ and neuro-endothelial interactions^50^, at least in part, regulate bone tissue healing via indirect effects on mesenchymal cell function.

These potentially important interaction effects require further isolated study. Finally, sex specific differences are well established in the incidence of stress fractures^51^ and in general the experience of orthopaedic pain^52^. Since there was no apparent sex difference in FGF9 expression in human DRG neurons^53^, we do not anticipate a sex difference even though all the animal studies were performed in male mice.

In sum, skeletal-innervating sensory neurons are a group of neurons with nociceptive and mechanoreceptive functions, which are transcriptionally responsive to bone injury in a temporal dynamic fashion, and positively regulate fracture healing via FGF9-FGFR signaling.

## Materials and Methods

### Animals

Mouse lines were used as previously described^1,2^, including TrkA^WT^ and TrkA^F592A^ mice (JAX: 022362) and C57BL/6 mice (JAX: 000664). TrkA^F592A^ mice were donated from the Ginty laboratory, which are homozygous for a phenylalanine-to-alanine point mutation in exon 12 of the mouse *Ntrk1* gene (F592A)^3^. This point mutation in TrkA^F592A^ mice renders the endogenous TrkA kinase sensitive to inhibition by the membrane-permeable small molecule 1NMPP1^3^. Adult male mice at 12-18 wk old were used for experiments unless otherwise indicated.

Littermate analysis was performed by investigators blinded to the experimental groups. Mice were kept under a 12-h light/dark cycle with food and water provided ad libitum. All procedures involving mice were approved by the Institutional Animal Care and Use Committee of The Johns Hopkins University under Protocol MO22M368.

### Retrograde labeling of skeletal-innervating neurons

In order to retrograde label peripheral sensory neurons, AAV-PHP.S-tdTomato (Addgene, 59462- PHP.S, titer ≥ 1×10¹³ vg/mL) was utilized, an engineered virus with enhanced tropism for peripheral neurons^4^. To validate the kinetics and specificity of retrograde tracing, a mixture of virus (3.5μl) and Fast Blue (1.5μl, Polysciences, Cat #:17740-1) was injected into the midshaft of the right ulnar periosteum. After 1-4 wks, ipsilateral and contralateral DRGs were harvested at levels Cervical7-8, Thoracic1 and Lumbar1.

### RNAscope

DRGs were fixed in 4% paraformaldehyde (PFA) overnight, sectioned at 14μm thickness and examined by RNAscope multiplex fluorescent reagent kit (Advanced Cell Diagnostics) per the manufacturer’s protocol using following probes: Mm-Asic3-O1 (Cat #:480541), Mm-Bmpr1b- C2 (Cat #:533941-C2), and tdTomato-C3 (Cat #:317041-C3).

### Whole mount immunohistochemistry

Whole mount immunohistochemistry of retrograde labeled DRGs was performed following the iDISCO protocol^5^. Briefly, DRGs were harvested and fixed using 4% PFA at 4°C overnight then subjected to serial methanol pretreatment. Tissues were then incubated with permeabilization solution and blocking solution for 1 d, and incubated with RFP Antibody pre-adsorbed rabbit polyclonal antibody in PBS/0.2% Tween-20 with 10 μg/ml heparin (PTwH)/5% DMSO/3% donkey serum for 3 d at 4°C. After washing with PTwH, DRGs were incubated with goat Anti-Rabbit IgG H&L and washed before proceeding to tissue clearing and imaging.

### *In vivo* interventions

#### Surgical transection

C57BL/6J WT mice were anesthetized with isoflurane gas (1-3%). After confirming deep anesthesia, a skin incision was made at the midpoint of the right upper arm to expose the ulnar nerve. A 3 mm segment of nerve distal to the ligature was excised. The incision was closed with a 5-0 Vicryl suture. A sham surgery was used, in which the nerve was exposed, but not excised. Validation of neurectomy was performed via examination of distal innervated skin at 1 wk post- operative.

#### Chemical TrkA inhibition

To inhibit TrkA catalytic activity, TrkA^WT^ and TrkA^F592A^ animals were used. 1NMPP1 (Aurora Analytics LLC) was administered to inhibit TrkA catalytic activity according to previously established protocol in TrkA^F592A^ animals^1^. 1NMPP1 has previously been shown to have no effect on fracture healing in TrkA^WT^ mice^1^. Briefly, 1NMPP1 was injected intraperitoneally 48 hr and 2 hr before loading using a 5nM solution, then maintained on drinking water at 40μM concentration until euthanasia.

### Induction of ulnar stress fracture

Stress fracture was produced via cyclic end-loading in 18-wk-old mice using previously established protocols^1,6^. Mice were anesthetized with inhaled isoflurane gas (1%-3%) during the period of the experiment. After confirming deep anesthesia, specially designed fixtures were used to secure the right olecranon process and the flexed carpus. Mechanical loading was performed using a material testing system (Instron, E1000 All-Electric Dynamic Test Instrument) that monitored force and displacement. Loading ended when a displacement constant was reached. Displaced fractures or combined ulnar/radial fractures were identified by high- resolution Faxitron imaging (Faxitron Bioptics), which occurred in a minority of mice and were excluded from analysis. A single subcutaneous injection of analgesic (1mg/kg buprenorphine) was provided after fracture.

### μCT imaging and analysis

Mice were euthanized14 d after fracture. Forelimbs were dissected and skin removed. Tissues were then fixed in 4% PFA for 24 hr at 4°C. SkyScan 1172 high-resolution μCT imaging system (Bruker) were used to acquire images (65 kV and 153μA with 1.0mm aluminum filter at 9μm resolution). NRecon (Bruker) were used for image reconstruction and CTAn software (Bruker) was used for quantification. Volumes of interest (VOI) of the callus region were used as previously described^1^, with measurements in accordance with the recommendations of the American Society for Bone and Mineral Research^7^.

### Single cell dissociation and sequencing

#### DRG tissue dissociation

Retrograde labeled whole DRGs (ipsilateral C7-T1 levels) were harvested at 0, 1, 14 and 56 d post-fracture. Dissection and dissociation were conducted using previously published protocols^8^, with minor modifications. Briefly, N=18 DRGs were isolated from 6 mice per timepoint into ice-cold PBS. DRGs were digested with 1.8ml digestion solution with Papain (25U/ml, Worthington, Cat #:LS003126), Collagenase A (20mg/ml, Sigma-Aldrich, Cat #:10103586001), Dispase II (20mg/ml, Roche, Cat #:61926600) and BSA (0.5%, Sigma-Aldrich, Cat #:A7979) at 37 °C.

30μl of Collagenase A/Dispase II (20mg/ml) was added depending on the dissociation progress. Once DRGs were dissociated, the cell suspension was filtered using a 40μm cell strainer (Corning), diluted with 3ml aCSF (Tocris Bioscience) and centrifuged at 100g for 10 min at 4°C. After removing the supernatant, the pellet was resuspended in 0.5ml aCSF and 0.5 ml complete Neurobasal medium. The cell suspension was then carefully layered on top of an Optiprep gradient (90μl Optiprep density solution (Sigma-Aldrich) + 455μl aCSF + 455μl complete Neurobasal). The gradient was centrifuged at 100 g at 4°C for 20 min. After centrifuging, the supernatant was removed until 100μl remained.

#### Callus tissue dissociation

Forelimbs were harvested 14 d after stress fracture (N=3 for each experimental condition). Callus tissue was dissected and marrow cells were flushed with PBS. After cutting into small pieces, fracture tissues were digested with 1mg/ml Collagenase 1/Collagenase 2 (Sigma-Aldrich) and 2 mg/ml dispase II (Roche). As soon as the cells were dissociated, cell suspension was centrifuged at 100g for 10 min at 4°C. The cell pellet was resuspended and incubated in ACK lysing buffer (Quality biological, Cat #:118156101) for 10 min. After centrifugation, cells were resuspended with PBS and filtered at 40μm (Corning). The dissociated cells were centrifuged again, and the supernatant was removed until 100μl remained.

#### Single cell RNA sequencing

The viability of dissociated cells was assessed with Trypan blue stain (Invitrogen, Cat #: T1-282) on a Countess II (ThermoFisher Scientific) automated counter and showed >90% viability in all cases. The cells were then sent to JHMI Transcriptomics and Deep Sequencing Core for single cell encapsulation, transcriptome capture, library preparation and sequencing. Libraries were generated using the 10X Genomics Chromium controller (Single Cell 3’ Dual Index Reagent Kits for DRGs and Single Cell 3’ HT Dual Index kit for callus tissue cells) following the manufacturer’s protocol, aiming for 10,000 cells per channel. Libraries were then sequenced and CellRanger 7.0.0 was used to perform sample de-multiplexing, barcode processing and single- cell gene counting. mm10-2020-A with tdTomato (NCBI reference number AQA28233.1) augmentation was used as genome reference for DRG cells and mm10-2020-A was used for callus cells. The above processes were achieved at the JHMI Transcriptomics and Deep Sequencing Core.

#### Bioinformatic analysis

Seurat ‘4.4.0’ ^9^ was used for downstream analysis. For neurons, cells with less than 1,000 genes expressed or with a fraction of mitochondrial gene UMIs higher than 0.075 were removed.

DoubletFinder was used to remove doublets^10^. Samples from different time points were integrated using the Harmony package^11^. For callus tissue, cells with less 200 genes expressed or with a fraction of mitochondrial gene UMIs higher than 0.15 were removed. Samples from different conditions were integrated with Harmony. Integrated Seurat objects were processed for clustering with standard Seurat 4 workflow. Clusters were labeled as cell types according to characteristic genes identified by Seurat FindMarkers function. An expression threshold of 0.3 was used to differentiate tdT^+^ and tdT^-^ neurons.

Pathway enrichment analysis and visualization in injured timepoints compared to uninjured neurons were performed by DAVID (https://david.ncifcrf.gov/home.jsp), Cytoscape (https://cytoscape.org/) and EnrichmentMap (https://apps.cytoscape.org/apps/enrichmentmap) according to published protocol with minor changes^12^. Briefly, DEG lists acquired from the comparisons of d1 vs uninjured, d14 vs uninjured and d56 vs uninjured were used as input for DAVID. Corresponding GO enrichment files were generated and were visualized in Cytoscape using EnrichmentMap. Single-cell regulatory network inference was achieved using SCENIC^13^ and visualized by Cytoscape. Temporal changes of gene expression corresponding to injury was analyzed and visualized by ClusterGVis (https://github.com/junjunlab/ClusterGVis). For GO enrichment in control callus compared with denervated callus, pseudobulk sequencing analysis was first performed to identify DEGs. GO enrichment and visualization were then achieved using clusterProfiler^14^ and enrichplot package (https://github.com/YuLab-SMU/enrichplot) for whole callus and DAVID for mesenchymal cells.

For interactome analysis, DRG neurons and fracture callus resident cells each from d14 post-fracture were used. Cellchat^15^ was utilized to identify interactions between neurons and fracture callus resident cells, mesenchymal cells and differential pathways between tdT^+/-^ neurons and mesenchymal cells with a minimum of 10 cells per cluster. In the computeCommunProb() function, the truncatedMean type-analysis was used to identify interactions between neurons and callus resident cells and the triMean type-analysis was used to produce interactions for differential pathways between tdT^+^ and tdT^-^ neurons with mesenchymal cells. Nichenet^16^ was used to predict active ligands secreted from tdT^+/-^ neurons that may interact with fracture callus mesenchymal cells.

### Human DRG collection and analysis

Human single-nuclei DRG data were integrated using Seurat’s CCA algorithm from three previously published studies. The collection of DRGs and data analysis processes can be found at Bhuiyan et al^17^.

### Histology and Immunohistochemistry of tissue sections

Tissues were harvested and placed in 4% PFA overnight at 4°C. For bone tissue, after washing in PBS for 1 hr, samples were decalcified in 14% EDTA (Sigma-Aldrich) for 21 d at 4°C. Tissues were then placed in 30% sucrose overnight at 4°C for cryoprotection before embedding in OCT and processed for cryosections at a thickness of 14 or 30μm. Thin sections were used for routine H&E and Safranin-O/Fast green staining. ALP staining was performed using prepared reagents following the manufacturer’s instructions (Sigma-Aldrich, Cat #:85L2-1KT). Histological images were taken using upright fluorescence microscopy (Leica Microsystems, Leica DM6). For immunofluorescent staining, thick sections were washed 3x with PBS for 5 min each. After permeabilization with 0.3% Triton-X for 20 min, sections were blocked with 5% normal goat serum for 1 hr and then incubated with primary antibodies (Supplementary Information File 10) overnight at 4°C. Sections were then washed 3x with PBS for 5 min each and incubated with appropriate secondary antibodies for 1 hr at RT. Finally, sections were mounted with Vectashield HardSet Antifade Mounting Medium with DAPI (Vector Laboratories, Vectashield H-1500) prior to imaging with confocal microscopy (Carl Zeiss Microscopy, Zeiss LSM800 FCS or LSM900 FCS). Images were analyzed by Imaris software. For periosteal nerve immunostaining quantification, five 20x three-dimensional volumetric regions of interest (200*1000 pixels) were centered on the ulna midshaft periosteum and analyzed. For IENF immunostaining quantification, five 20x three-dimensional volumetric regions of interest (100*100 pixels) aligned with epidermis were analyzed. Proliferation and osteogenesis were measured by Ki67 and OCN fluorescence intensity, both quantified using whole sections of fracture callus tissue using Fiji software^18^.

### *In vitro* experiments

#### Cell culture

Mouse DRG neuronal culture was performed using previously established protocols^1,2^. DRGs were harvested from 18-wk-old male C57BL/6J mice. After digestion as above, dissociated neurons were plated into 24-well plates coated with 100μg/ml poly-D-lysine (Sigma-Aldrich) and 10μg/ml laminin (Gibco). Neural basal media with anti-mitotics included 5% FBS, 1% penicillin/streptomycin (Gibco), 1X Glutamax (Gibco), 2% B27 (Thermo Fisher Scientific), 20 μM 5-fluoro-2-deoxyuridine and 20 μM uridine (Sigma-Aldrich).

Isolation of mouse periosteal cells was conducted as described^19^. Briefly, hindlimbs were dissected, and the surrounding soft tissues were carefully removed to expose the tibial and femoral periosteum (N=6 C57BL/6J 18-wk-old mice per cell isolation). The tissue was then washed twice in PBS with antibiotic and antimycotic and digested with 2mg/ml collagenase P (Roche) in DMEM (Gibco) containing 0.5% BSA (Sigma-Aldrich) at 37°C for 10 min (6 times). The first two digestions were discarded, and the rest of the digestions were pooled. The digestion was filtered with 40μm cell strainer and centrifuged at 2,000rpm for 20 min, resuspended and cultured in αMEM (Gibco) with 20% FBS (Gibco), 1% penicillin/streptomycin. Medium was changed every 72 hr. All *in vitro* experiments were performed at passage 2, with biologic and technical triplicates performed.

#### siRNA knockdown

DRG neuron electroporation was performed as previously described^2,20^ with minor modifications. DRGs were harvested and digested as described above. The dissociated cells were then filtered with 40 μm cell strainer and centrifuged at 100 g for 10 min. The pelleted cells were resuspended with 100μl electroporation buffer (Lonza, Cat #: VPG-1001) containing 0.3 nmol nontargeting control siRNA (siRNA scramble, Scr) or siRNA against *Fgf9* mRNA (Locus ID 14180, Cat #:SR405443) and transferred to a 2.0 mm electroporation cuvette and electroporated with Nucleofector II (Lonza) following the manufacturer’s manual. After electroporation, the cells were mixed with 500μl of Neural basal media with 5% FBS, 1% penicillin/streptomycin, 1X Glutamax, 2% B27 supplement and anti-mitotic reagents and seeded into precoated 24-well plates. Media was changed the following morning. Knockdown efficiency was confirmed with quantitative real-time polymerase chain reaction (qRT-PCR) on d 4.

#### Neuronal Conditioned media (CM) collection

To collect neuronal CM, DRG neurons were cultured for 4 d to establish axonal networks in neural basal media. The media was then changed to αMEM with 1% FBS and 1% penicillin/streptomycin (Gibco) on d4. From this point, the CM was harvested every d for 5 d. CM was filtered with 0.22 µm cell strainer was stored at -80 °C until further use.

#### MTS proliferation assay

Periosteal cells at passage 2 were cultured in αMEM, 10% FBS until 80% confluence was reached (5,000 cells per well in 96-well plates). In select experiments, periosteal cells were treated either with neural CM, PD173074 (100nM, Sigma-Aldrich), or recombinant FGF9 (5-25 ng/ml, R&D Systems, INC, Minneapolis, MN). After 24, 48, and 72 hr MTS assay was performed following the manufacturer’s instructions (Promega).

#### Osteogenic differentiation

Periosteal cells at passage 2 were seeded at a density of 2 x 10^5^ per well in 24-well plates for osteogenic induction (or 48-well plates for *Fgf9* KD experiment). Osteogenic media included 10% FBS, 1% Penicillin/streptomycin with 10mmol/L β-glycerolphosphate, 50µmol/L of Ascorbic acid and 1mmol/L of dexamethasone. In select experiments, medium included rmFGF9, PD173074, neuronal CM, or neuronal CM with *Fgf9* knockdown. Media was changed every other d until 28 d. At the end point of differentiation, cells were washed and fixed with Alizarin red staining (ARS) and quantified as per our previously published protocol^21^.

Osteogenic gene expression was assessed by qRT-PCR.

#### qRT-PCR

GeneElute single cell RNA isolation kit (Sigma Aldrich, USA) was used for total RNA isolation. Subsequently, 1μg of RNA was used to synthesize cDNA synthesis using an iScript cDNA Synthesis Kit (Bio–Rad, Hercules, CA). Real-time PCR was performed using SYBR™ Green PCR Master Mix (Life Technology), and detection was performed with a QuantStudio 5 Real-Time PCR system instrument (Thermo Scientific, Waltham, MA). GAPDH was used as an internal control for all genes. Primer information is provided in **Supplementary Information File 12**.

### Somascan assay of human DRGs and sciatic nerves

The study was approved by The Institutional Review Boards of UT Dallas (UTD) and MD Anderson Cancer Center (MDACC). All experiments conformed to relevant guidelines and regulations. Dorsal root ganglion tissues were collected from the donors after neurological determination of death within 2-3 hours of cross-clamp (Donor information can be found in **Supplementary Information File 13**). Tissue lysates were prepared from fresh-frozen DRG. The tissues were placed in T-PER Tissue Protein Extraction Reagent (Thermo Scientific, Cat # 78510) with additional 1X Halt Protease Inhibitor Cocktail (Thermo Scientific, Cat # 87786) and homogenized using Precellys Soft Tissue Homogenizing beads (Bertin Corp, Cat # P000933-LYSK0-A.0). Samples were centrifuged at 14,000 x g for 15 minutes in the cold room. The resulting supernatant was quantified by Micro BCA™ Protein Assay Kit (Thermo Scientific, Cat# 23235) and normalized accordingly. Proteins were profiled using the SOMAScan platform (SomaLogic, Inc., Boulder, CO). 7000 analytes were measured on the SOMAScan assay. Quality controls were performed by SomaLogic to correct for technical variabilities within and between runs for each sample^22^.

### Statistics

All statistical analyses were performed using Prism (GraphPad). A Shapiro-Wilk test was used to test normality for all datasets. Parametric data was analyzed using Student’s *t*-test for two-group comparison, or a one-way ANOVA for multi-group comparison, followed by Tukey’s multiple comparisons test. Quantitative data are expressed as mean ± 1 SD, with \**p<*0.05, \*\**p<*0.01 considered statistically significant. Power analyses were performed to determine sample size. A one-tail test with effect size of d=1.5 for denervation experiments (based on our previous published work^1^) and significance level of α=0.2 was used, suggesting a sample size of n=7 per group using G*Power software.

## Acknowledgement

We thank Drs. Jihye Yea, Soohyun Kim, Ziyi Wang, and Ray Cheng and the JHU microscopy facility and JHMI Transcriptomics and Deep Sequencing core for their technical assistance. AWJ was supported by the following funding sources: NIH/NIAMS (P01 AG066603, R01 AR079171, R21AR078919), NIH/NIDCR (R01 DE031488, R01 DE031028), Alex’s Lemonade Stand Foundation (22-26743), American Cancer Society (DBG-23-1155131-01-IBCD), the Maryland Stem Cell Research Foundation (2021-MSCRFD-5641), and Department of Defense (USAMRAA HT9425-24-1-0051). The content is solely the responsibility of the authors and does not necessarily represent the official views of the National Institute of Health, Department of Defense, or U.S. Army.

## Author contributions

MX, TLC and AWJ conceptualized the project. MX, NT, MC, ZL, MASG, CZ, XWW, RT, PC, YG developed the methodology. MX, NT, MC, ZL, MYZ, XX, CZ, XWW, JM and IS acquired and analyzed data. MX, ZL and MC performed animal studies. MX, ZL, DTF and MASG performed computational analyses. AWJ and TP provided transcriptomic data. AWJ and TLC provided transgenic mice. AWJ supervised the project and secured funding. MX and AWJ were responsible for writing the original draft. AWJ handled review and editing of the manuscript.

## Additional Information

Supplementary information is available. Correspondence and requests for materials should be addressed to A.W.J.

## Supplemental Information

**Supplemental Information File 1**: List of significantly increased secreted genes in tdT^+^ neurons compared to tdT^-^ neurons by scRNA-Seq.

**Supplemental Information File 2:** Significantly enriched GO terms in injured conditions (d1, d14, d56) when compared to uninjured condition by DAVID in tdT+ neurons by scRNA-Seq.

**Supplemental Information File 3:** Gene list of each category by Mfuzz clustering in tdT^+^ neurons tdT^-^ neurons by scRNA-Seq.

**Supplemental Information File 4:** Significantly enriched GO terms in injured conditions (d1, d14, d56) when compared to uninjured condition by DAVID in tdT^-^ neurons by scRNA-Seq.

**Supplemental Information File 5:** Significantly differentially expressed genes (|Log2FC| > 1 and Padj < 0.05) from pseudobulk sequencing analysis of all callus cells among control vs denervation groups by scRNA-Seq.

**Supplemental Information File 6:** Significantly enriched GO terms in denervation group when compared to control group by DAVID among all callus cells by scRNA-Seq.

**Supplemental Information File 7:** Significantly differentially expressed genes (|Log2FC| > 1 and Padj < 0.05) from pseudobulk sequencing analysis among callus mesenchymal cells from control vs denervation groups by scRNA-Seq.

**Supplemental Information File 8:** Significantly enriched GO terms among callus mesenchymal cells from control as compared to denervation groups by DAVID using scRNA-Seq.

**Supplemental Information File 9:** Significantly enriched GO terms among callus mesenchymal cells from denervation as compared to control groups by DAVID using scRNA-Seq.

**Supplemental Information File 10:** Gene lists of module scores used.

**Supplemental Information File 11:** Antibodies used.

**Supplemental Information File 12:** Primers used.

**Supplemental Information File 13:** Donor information of somascan assay.

## Data Availability and Code Availability

Requests for data and materials should be addressed to A.W.J. scRNA-seq data of mouse are deposited in GEO (GSE265817). Human DRG single cell sequencing data are available at https://harmonized.painseq.com/

## Extended Data Figures and Figure legends

**Extended Data Fig. 1.**
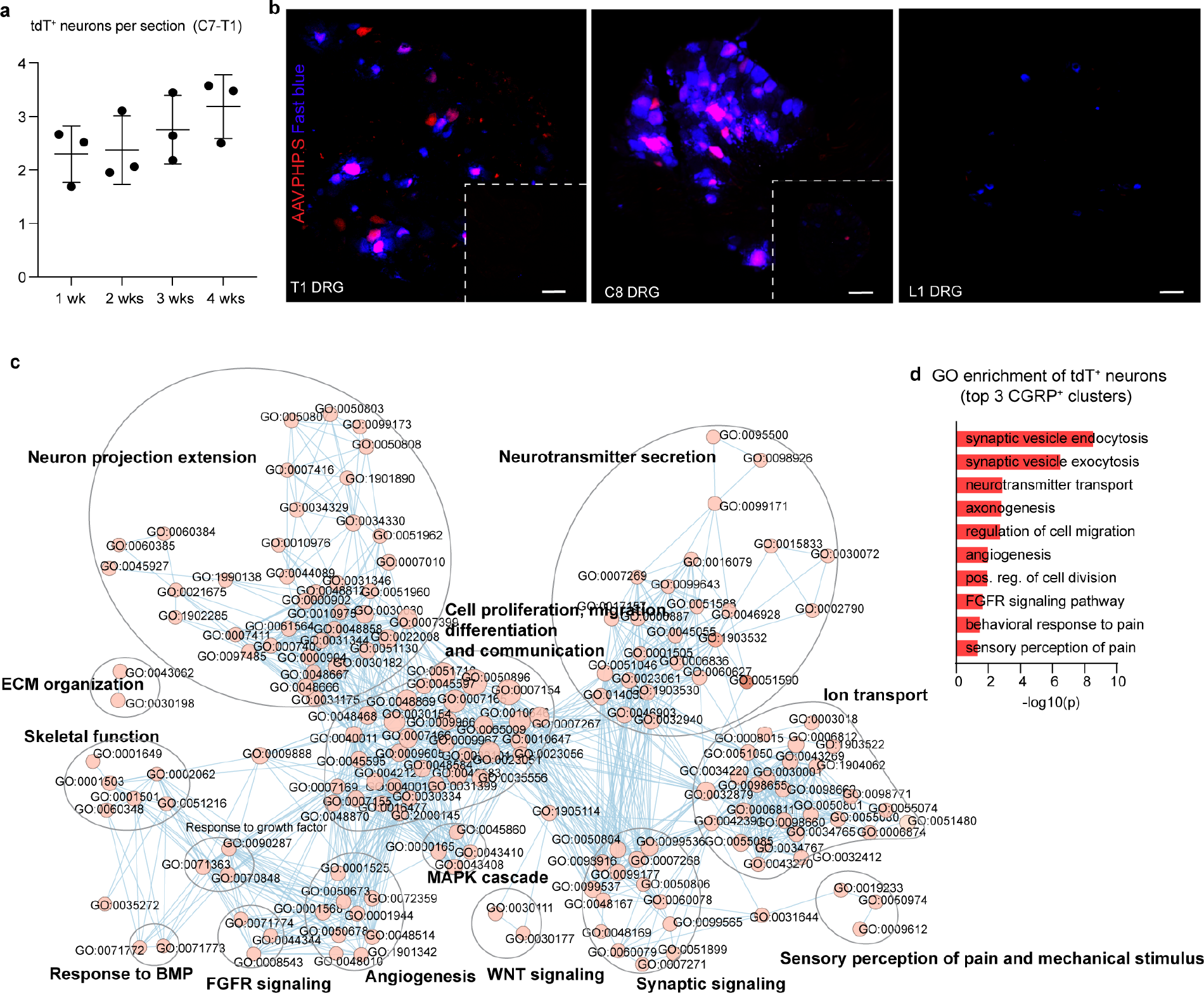
AAV-tdT labeling kinetics, specificity for DRG neurons, and GO term enrichment in tdT^+^ neurons. **a** tdT^+^ neuron frequency per section from ipsilateral C7-T1 at 1-4 wks post-injection of AAV-PHP.S into the ulnar mid-diaphyseal periosteum. N=3 mice (9 DRGs) per timepoint. **b** Co-visualization of tdTomato and fast blue in ipsilateral T1, C8 and L1 DRGs. Images correspond to Fig. 1b. Inset shows contralateral DRGs at the same spinal cord level. **c** GO term enrichment in tdT^+^ neurons (complete labeled version corresponding to Fig. 1i). Enrichment map displays significantly enriched gene ontology (GO) in tdT^+^ neurons as compared to tdT^-^ neurons. Nodes represent gene-sets and edges represent GO defined relations. Clusters are annotated according to the corresponding function. **d** Curated GO terms enriched in tdT^+^ neurons as compared to tdT^-^ neurons in top three neuron clusters with highest AAV-tdT labeling (CGRP-β/γ, CGRP-ζ and CGRP-η). Data obtained from N=1,020 neurons, N=12 DRGs (C7, C8 and T1) from N=4 mice. Scale bar: 100μm.

**Extended Data Fig. 2.**
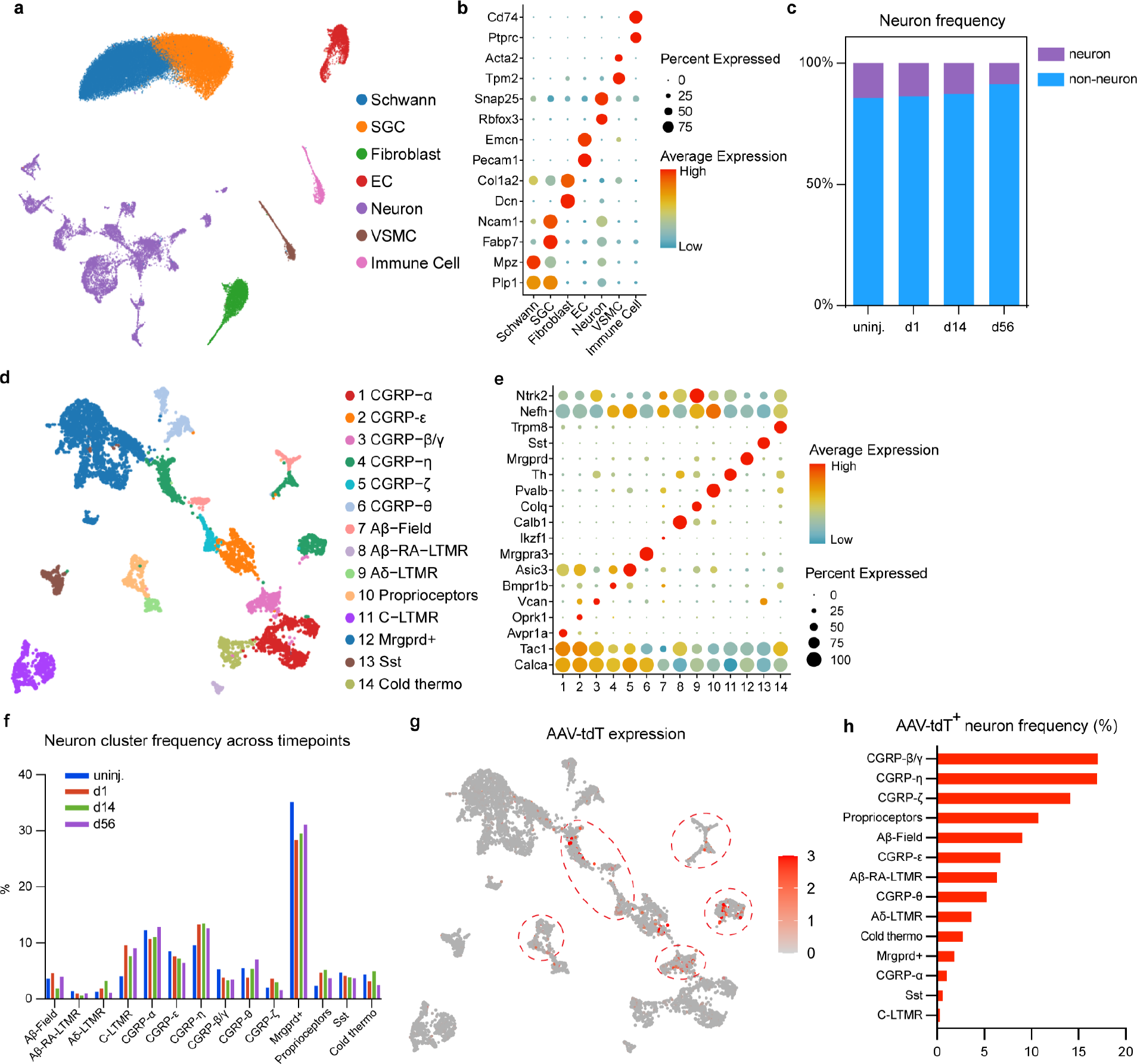
Cell composition and AAV-tdT labeling across timepoints after bone injury. Mice were subjected to mid-diaphyseal periosteal labeling by AAV.PHP.S 4 wks prior to stress fracture. Ulnar stress fracture was performed on 18-wks-old C57BL/6J mice. Ipsilateral C7-T1 DRGs were harvested for scRNA-seq at the uninjured time point, as well as 1, 14, and 56 d post-fracture. **a** UMAP visualization of all DRG cells by scRNA-seq obtained from all timepoints combined. SGC: Satellite glial cell; EC: Endothelial cell; VSMC: Vascular smooth muscle cell. **b** Dot plot of representative marker gene expression for each DRG cell type. **c** Neuron cell frequency at each timepoint. **d** UMAP visualization of all neuronal cells, obtained from all timepoints combined. **e** Dot plot of representative marker gene expression for each neuron cluster. **f** Percentages of each neuron clusters across timepoints. **g** UMAP of neuronal AAV labeling identified by *AAV-gene1,* all timepoints combined. **h** Frequencies of AAV-tdT labeling of neuron clusters, all timepoints combined. N=12 DRGs from 4 mice were obtained for the uninjured timepoint. N=18 DRGs from 6 mice were obtained per timepoint post-fracture. N= 53,553 total cells and N= 6,648 total neurons obtained.

**Extended Data Fig. 3.**
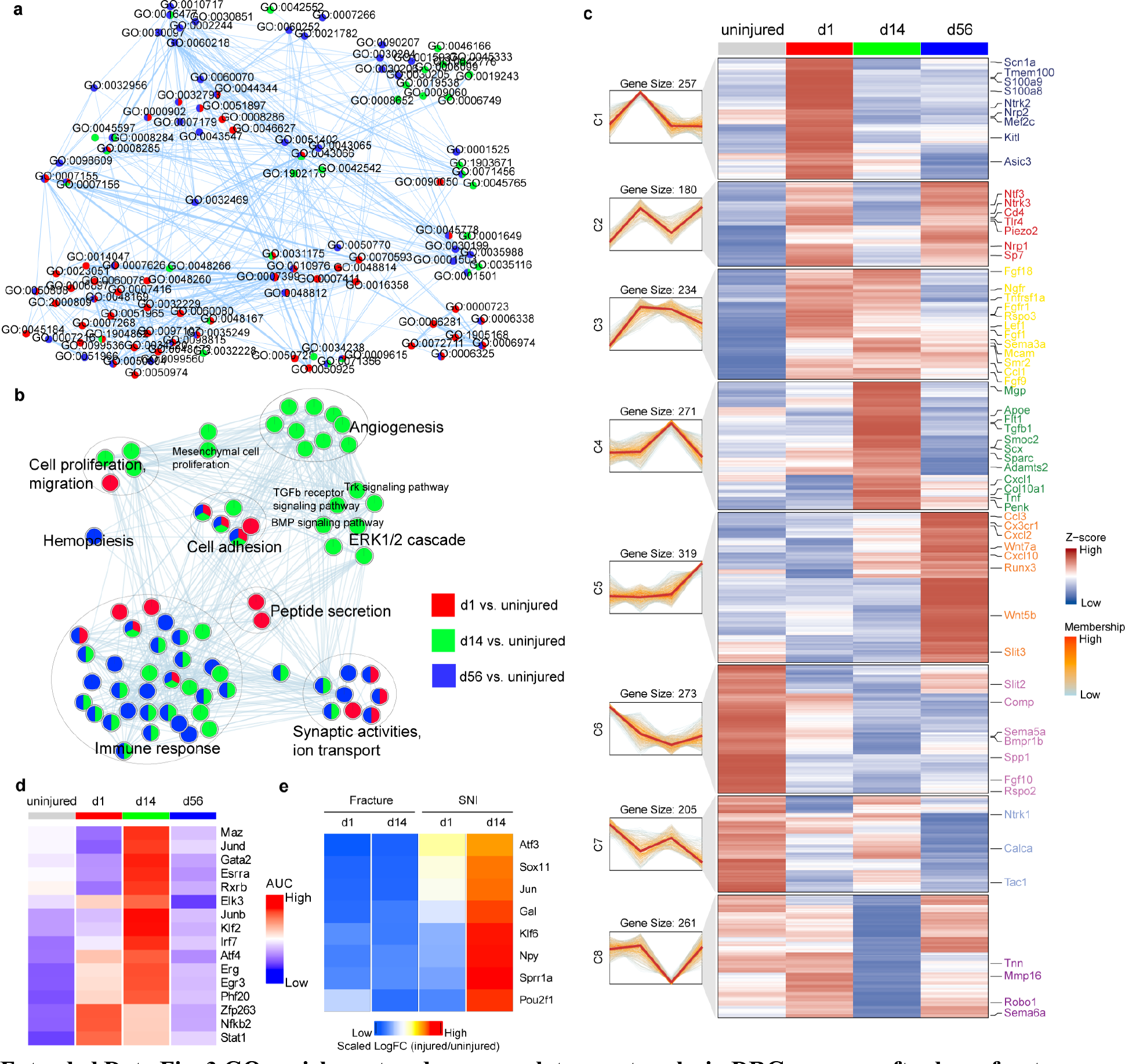
GO enrichment and gene regulatory networks in DRG neurons after bone fracture. A. GO enrichment map in tdT+ neurons after bone fracture (complete labeled version corresponding to Fig. 2b). Enrichment map displays the significantly enriched gene-sets in neurons after fracture vs. uninjured condition. Nodes represent gene-sets and edges represent GO defined relations. Clusters are annotated according to the corresponding function. N=12 DRGs from 4 mice were obtained for the uninjured timepoint. N=18 DRGs from 6 mice were obtained per timepoint post-fracture. N=389 total neurons analyzed. **b** GO enrichment map in tdT- neurons after bone fracture. Enrichment map displays the significantly enriched gene-sets in neurons after fracture vs. uninjured condition. Nodes represent gene-sets and edges represent GO defined relations. Clusters are annotated according to the corresponding function. N=6,259 total neurons analyzed. A full list of GO terms can be found in **Supplemental Information File 4**. **c** Soft clustering of 2,000 highly variable genes across four different timepoints by Mfuzz using ClusterGvis in tdT- DRG neurons. N= 6,259 neurons in total. **d** Heatmap of significantly increased transcription factor (TF) activity in tdT+ neurons 1, 14 and 56 d after injury in comparison to the uninjurd timepoint by SCENIC. **e** Heatmap of relative expression of common neuronal markers of nerve injury at d1 and d14 compared to uninjured condition in the fracture model versus a previously published spared nerve injury (SNI) model^21^.

**Extended Data Fig. 4.**
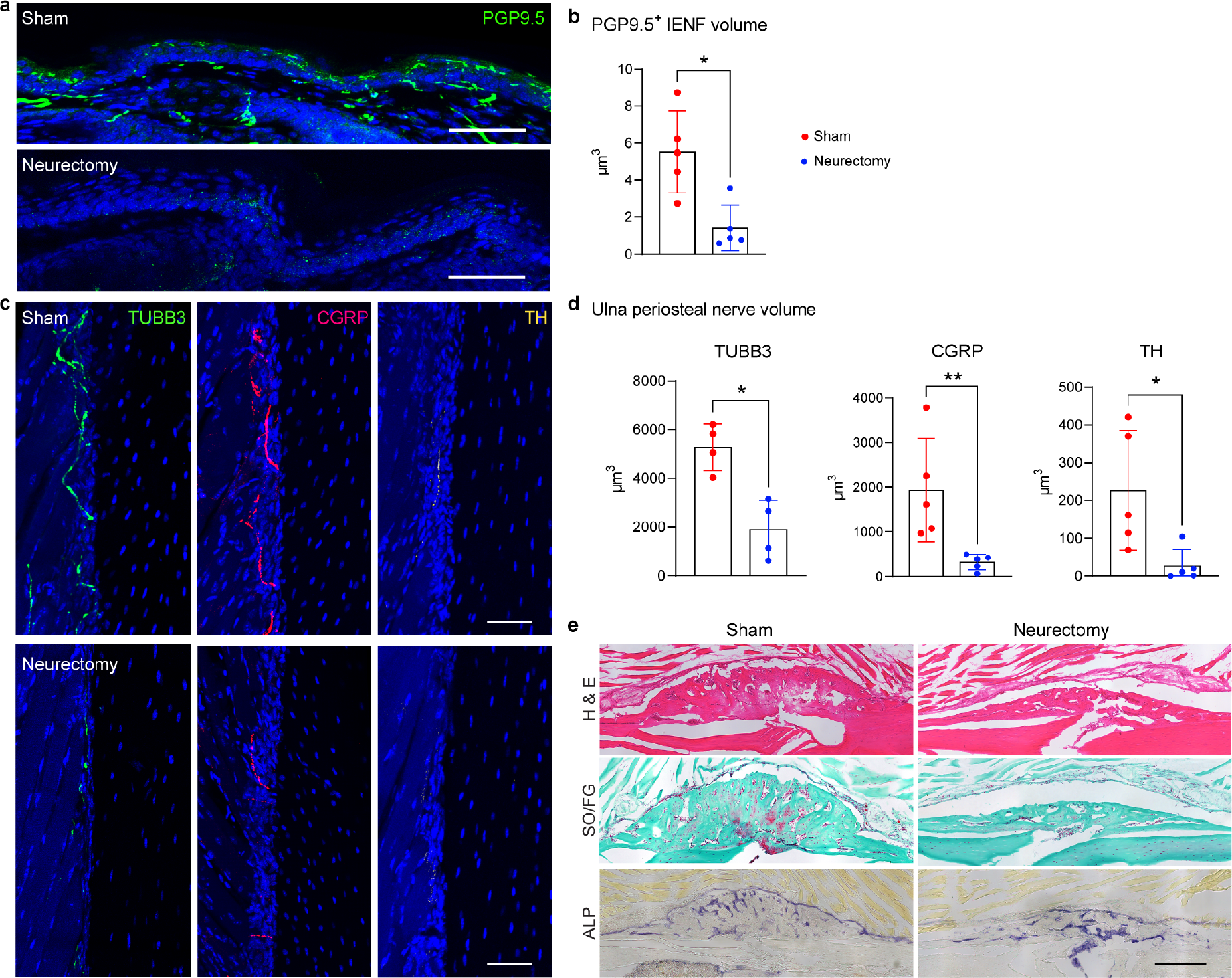
Ulnar nerve neurectomy results in target tissue denervation and compromised callus formation. a-d. C57BL/6J male mice underwent ulnar nerve neurectomy or sham surgery, followed by analysis one wk thereafter. **a,b** Intraepidermal nerve fiber frequency (IENF), shown by representative images of PGP9.5 immunostaining and quantification of skin of the 5^th^ digit, 1 wk after neurectomy or sham surgery. **c** Representative images of pan-neuronal marker TUBB3 (Beta III Tubulin), sensory neuronal marker CGRP (calcitonin gene related peptide) and sympathetic neuronal marker TH (Tyrosine hydroxylase) immunostaining of midshaft ulna periosteum, 1 wk after neurectomy or sham surgery. **d** Quantification of TUBB3^+^, CGRP^+^ and TH^+^ nerve fibers of the midshaft ulna periosteum. Scale bar: 50μm. N=5 mice per group. Values plotted are the means with errors bars representing ± 1 SD. **e** Histology of fracture callus from sham and neurectomy groups. C57BL/6J male mice underwent ulnar nerve neurectomy or sham surgery, followed by end-loading induced stress fracture of the ulna 1 wk thereafter. Histologic analysis performed 2 wks post-fracture. Representative H&E, Safranin O/Fast Green (SO/FG) and alkaline phosphatase (ALP) staining from sham and neurectomy groups. Scale bar: 250μm. N=6 mice per group.

**Extended Data Fig. 5.**
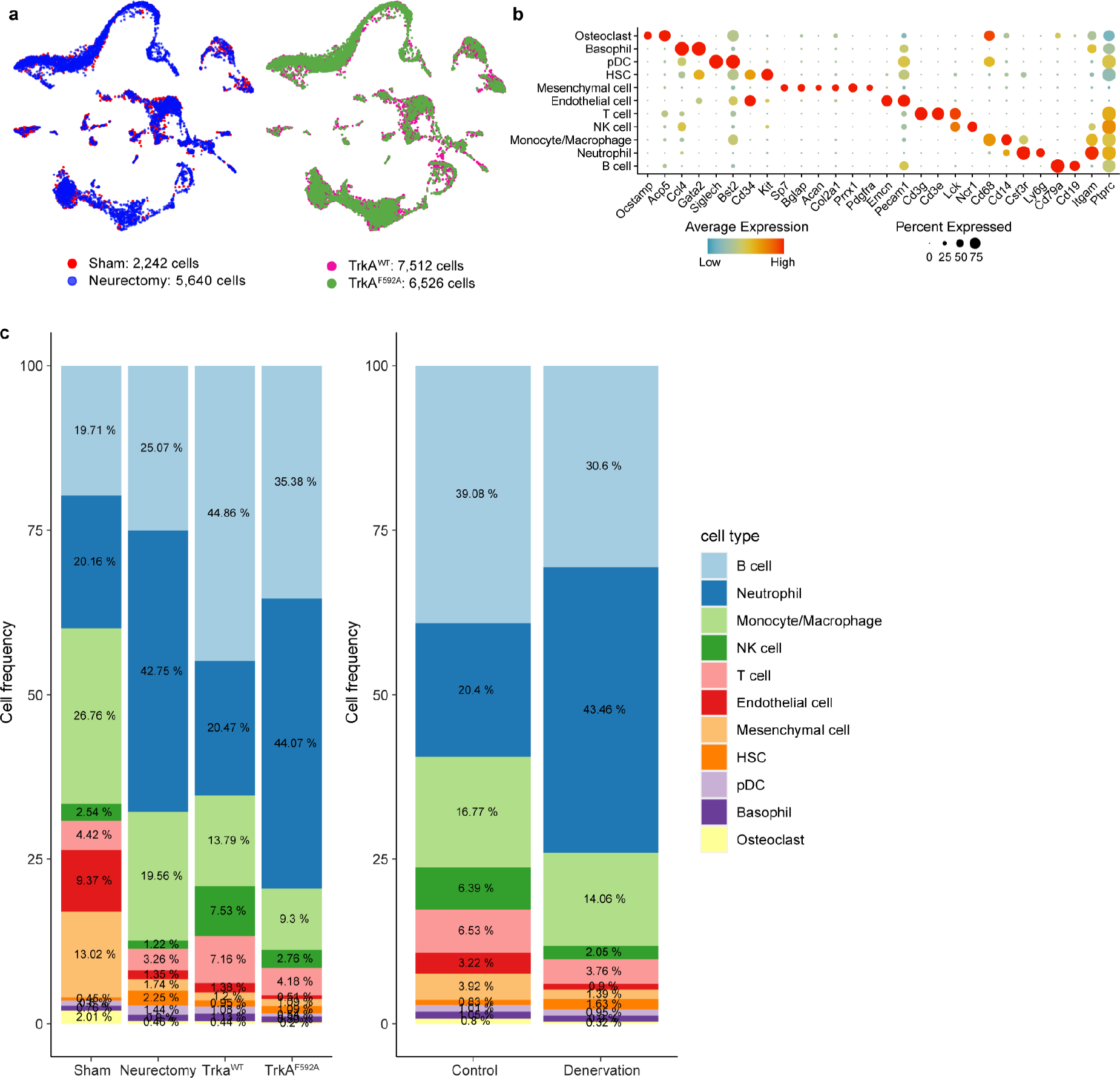
Cell composition of the fracture callus by scRNA-seq across experimental conditions. Two parallel experiments to denervate the fracture site were performed, via either surgical or chemical-genetic denervation. Surgical denervation was achieved via ulnar nerve transection, sham surgery was performed with visualization but not transection of the nerve as a control. Chemical-genetic denervation was achieved by 1NMPP1-treated TrkA^F592A^ mice with 1NMPP1-treated TrkA^WT^ mice as control. Stress fracture was performed on mice from both experiments at 18 wks old and fracture callus tissue was harvested at d14 post-fracture for scRNA-seq. **a** UMAP visualization of fracture callus scRNA- seq data with total cell numbers shown. **b** Dot plot of marker gene expression in each cell cluster. **c** Percentages of each cell type in each experimental condition (left). Also shown are cell percentages across ‘control’ (Sham and TrkA^WT^ groups) or ‘denervation’ conditions (Neurectomy and TrkA^F592A^ groups) (right). N=21,920 total cells, N=3 mice per group.

**Extended Data Fig. 6.**
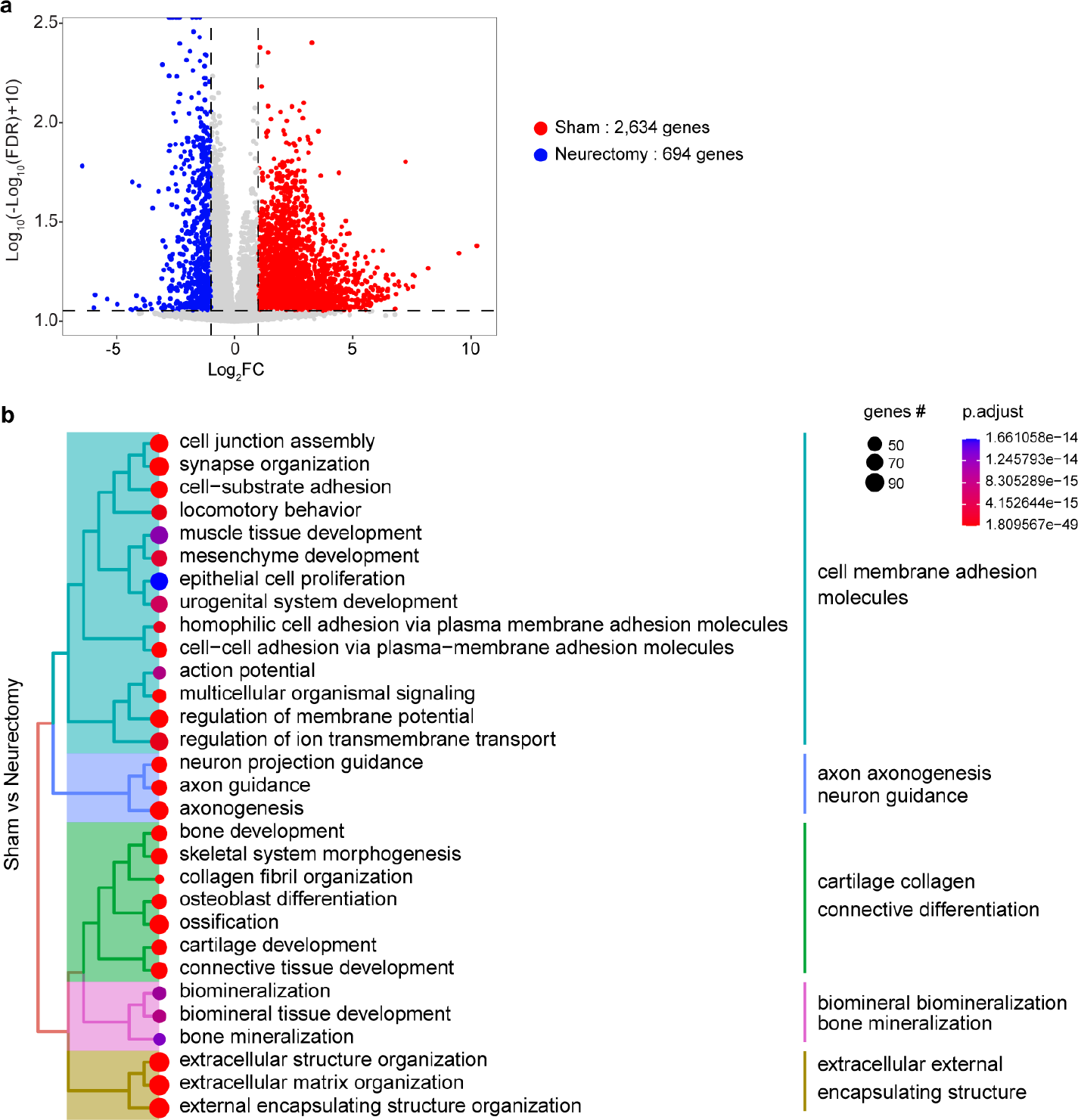
Additional scRNA-seq analyses of fracture callus tissue in the model of surgical denervation. Surgical denervation was achieved via ulnar nerve transection, sham surgery was performed as a control. One wk thereafter, stress fracture was performed on mice from both experiments at 18-wks of age, and fracture callus tissue was harvested at d14 post-fracture for scRNA-seq. **a** Volcano plot of overrepresented genes in sham and neurectomy groups among all cell clusters. Significantly differentially expressed genes (DEGs) identified using thresholds with FDR < 0.05 and Log2FC > 1. Total DEGs shown. **b** Tree plots of GO term enrichment in all cell clusters in sham group in comparison to neurectomy group. N=3 mice per group.

**Extended Data Fig. 7.**
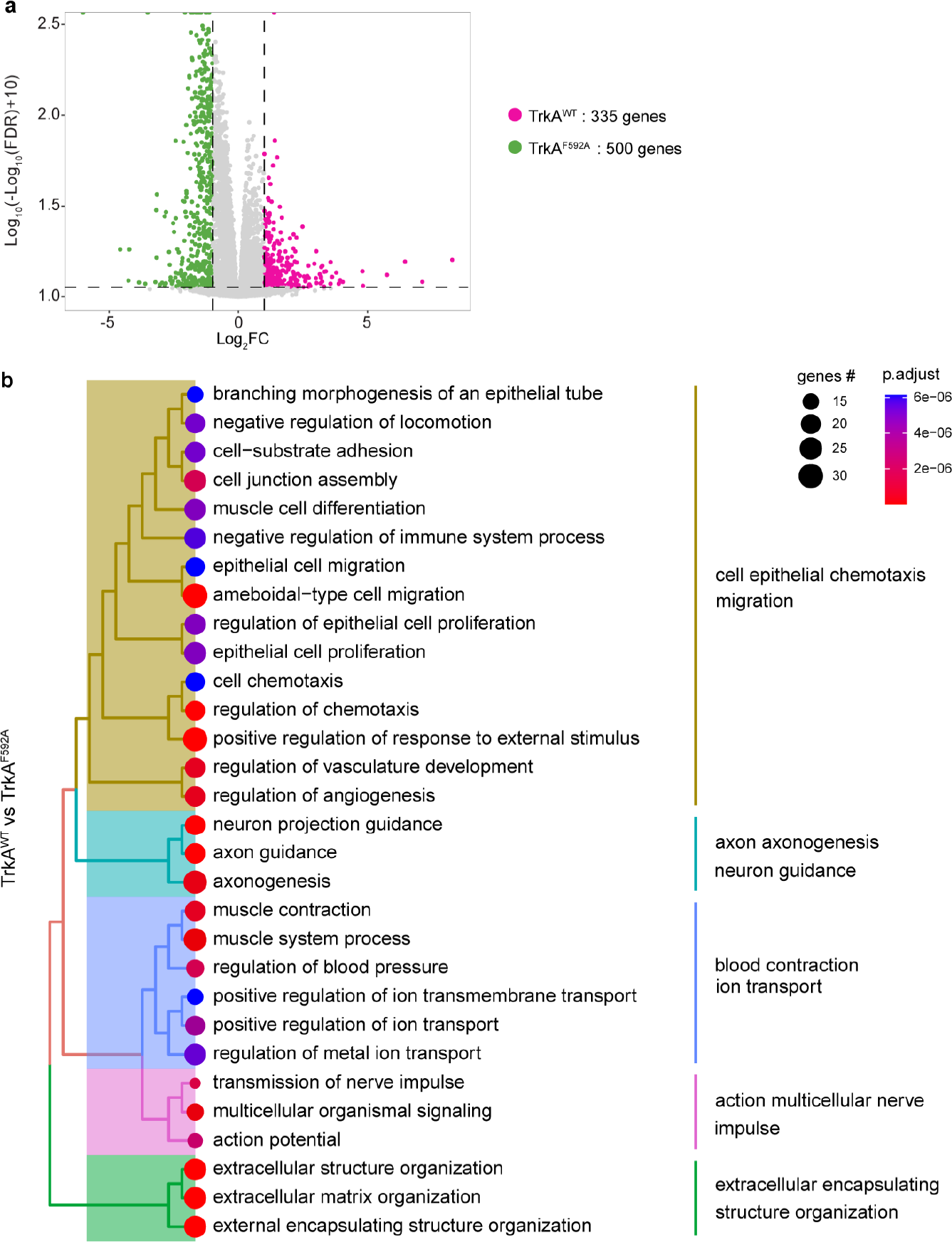
Additional scRNA-seq analyses of fracture callus tissue in the model of chemical-genetic denervation. Chemical-genetic denervation was achieved by 1NMPP1- treatment of TrkA^F592A^ mice with 1NMPP1-treated TrkA^WT^ mice used as a comparison. Stress fracture was performed on mice from both experiments at 18-wks of life, and fracture callus tissue was harvested at d14 post-fracture for scRNA-seq. **a** Volcano plot of overrepresented genes in TrkA^WT^ and TrkA^F592A^ groups in all cell clusters. Significantly differentially expressed genes (DEGs) are identified using thresholds with FDR < 0.05 and Log2FC > 1. Total DEGs shown. **b** Tree plots of GO term enrichment in all cell clusters in TrkA^WT^ group in comparison to TrkA^F592A^ group. N=3 mice per group.

**Extended Data Fig. 8.**
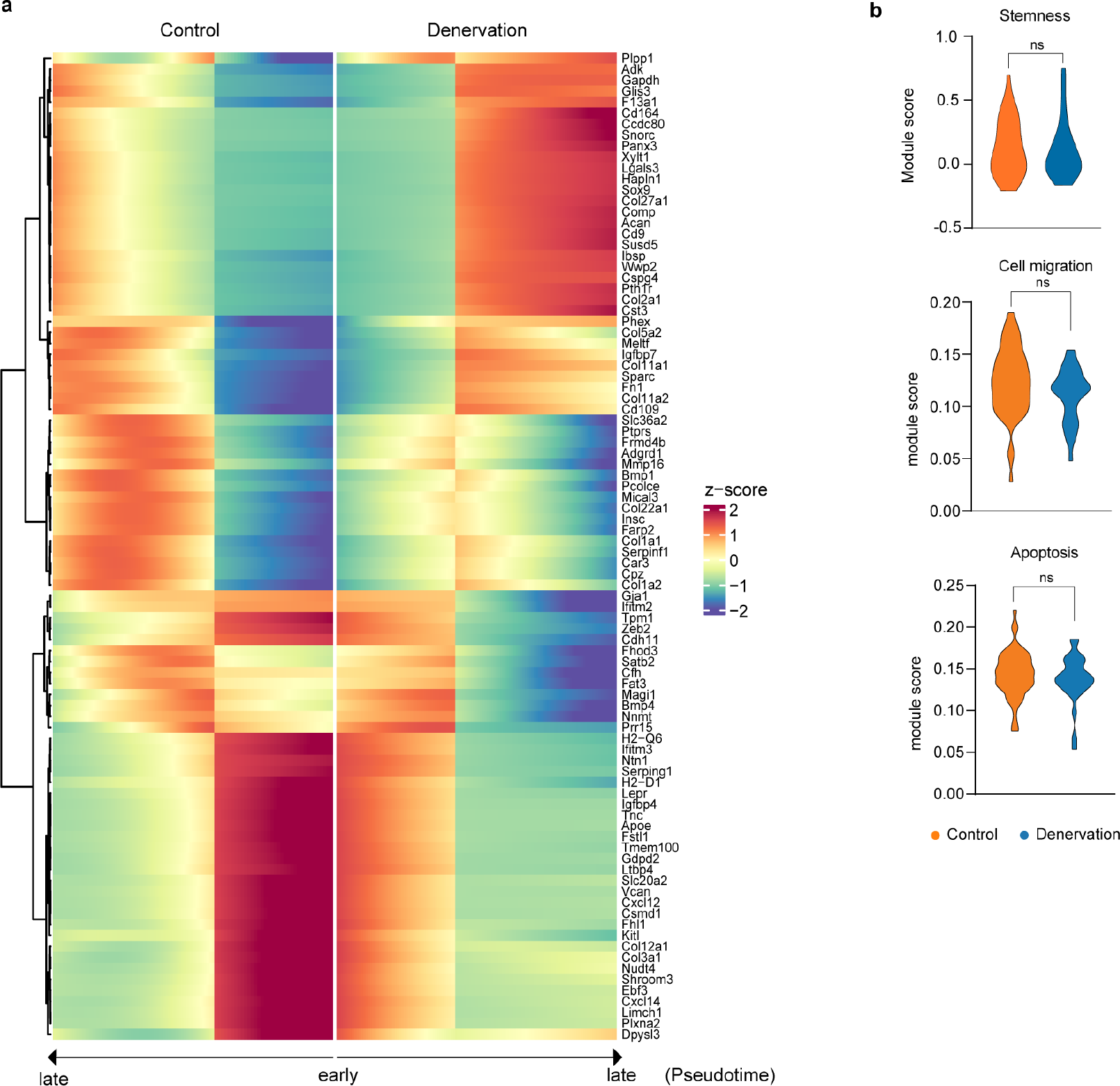
Pseudotime and additional module score analyses in fracture callus scRNA-seq. Sequencing analysis performed at 14d post-fracture among control or denervated callus tissue. Mesenchymal cells only presented for analysis. **a** Heatmap of genes by hierarchical clustering across pseudotime in both control and denervation groups. **b** Module score analysis performed within the *Pdgfra*^+^*Lepr*^+^ stem-like cells. Violin plots of module scores shown. Gene lists in module scores shown in Supplemental Information File 9. ns: non-significant.

**Extended Data Fig. 9.**
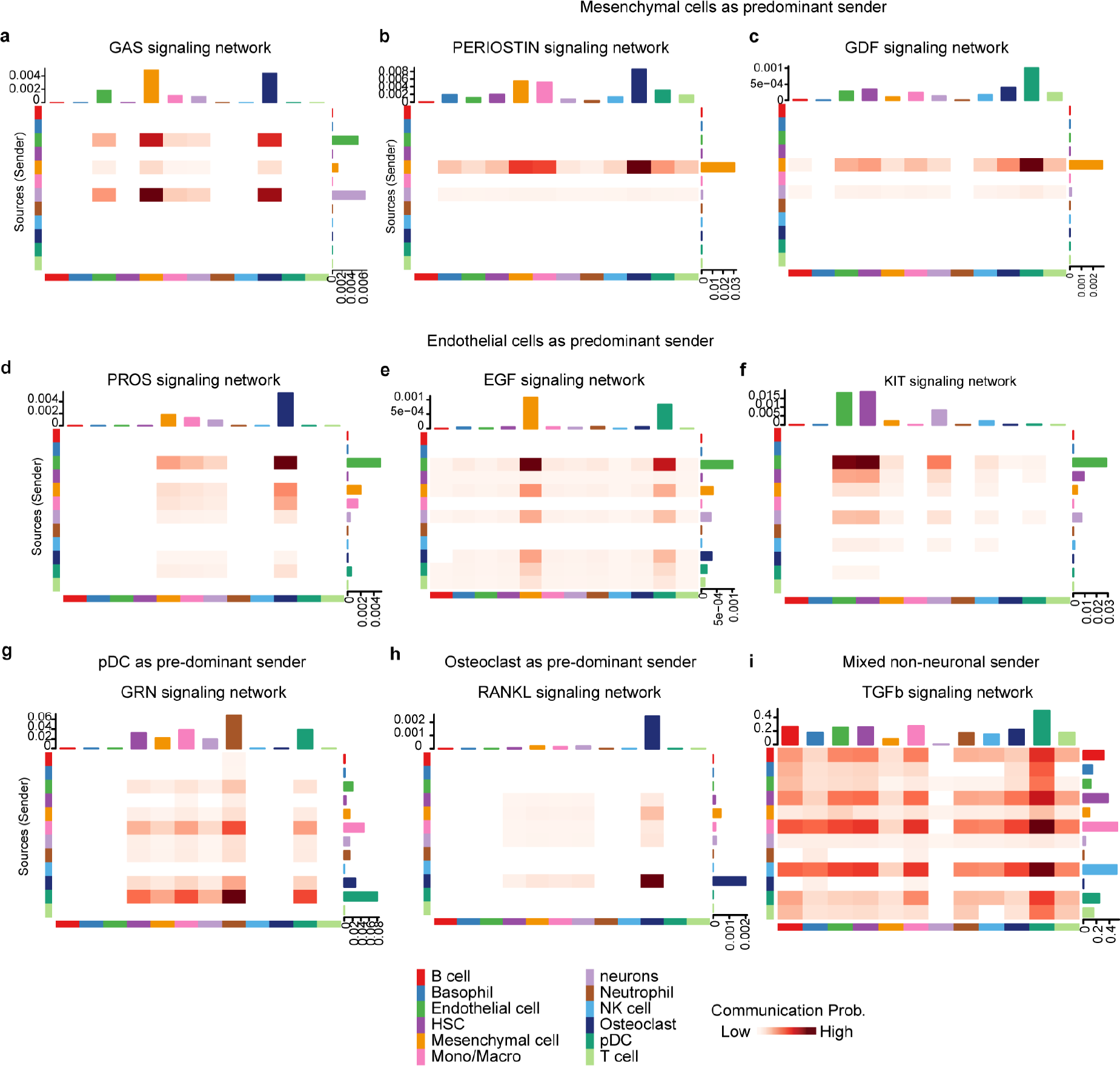
Heatmaps of interactions of signaling pathways network from senders to receivers. Combined scRNA-Seq datasets of DRG neurons and callus tissue, both derived from d14 post-fracture, and analyzed using CellChat. Shown are signaling pathways which did not demonstrate predicated neuron-to-callus signaling. **a**, Signaling pathways with neuron cells as the predominant sender cell (GAS signaling), **b, c** Signaling pathways with mesenchymal cells as the predominant sender cell (Periostin and GDF signaling), **d-f** Signaling pathways with endothelial cells as the predominant sender cell (PROS, EGF and KIT signaling), **g** Signaling pathways with pDC as the predominant sender cell (GRN signaling), **h** Signaling pathways with osteoclasts as the predominant sender cell (RANKL signaling), and **i** mixed non- neuronal sender cells (TGFb signaling).

**Extended Data Fig. 10.**
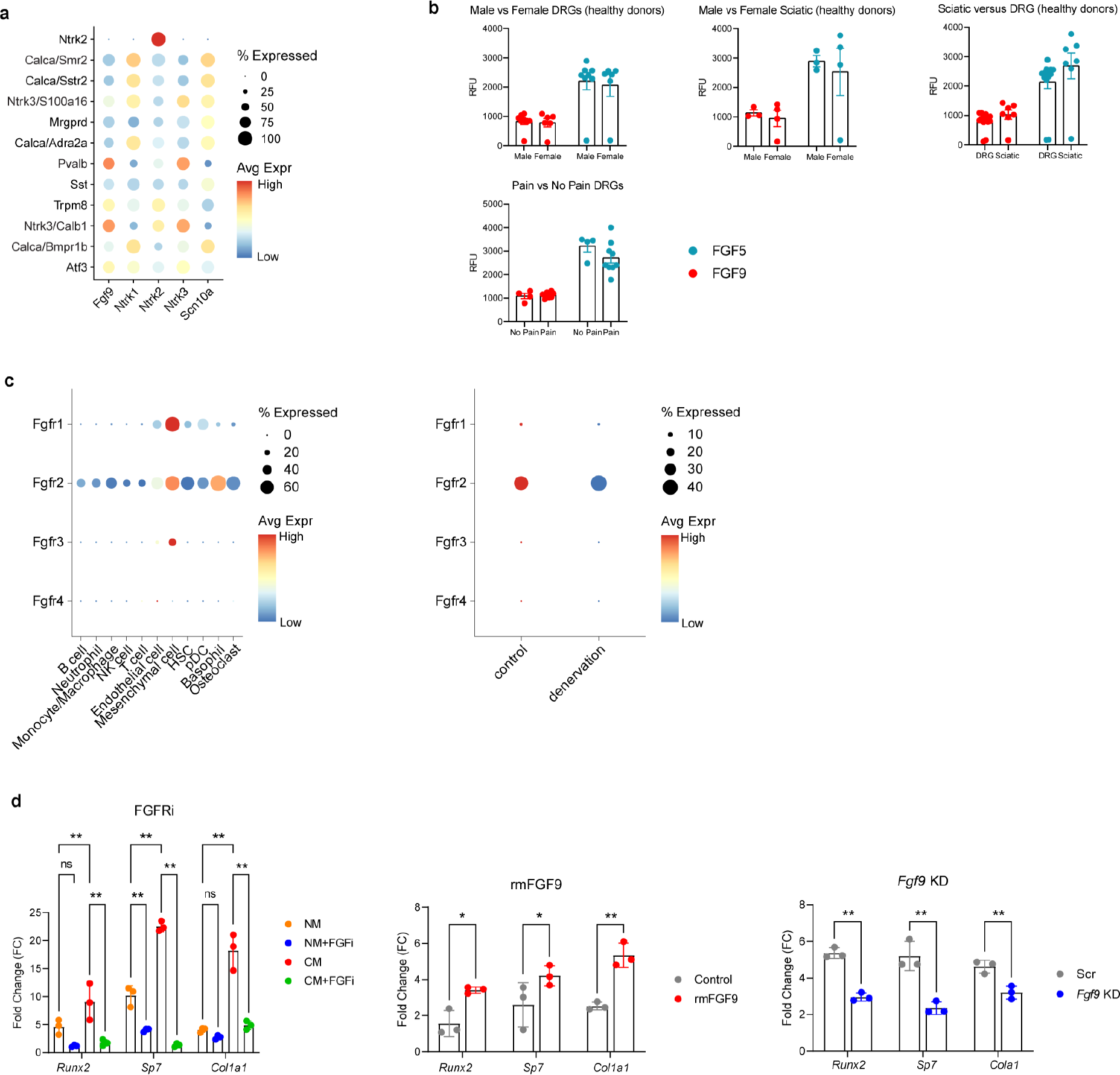
Expression of fibroblast growth factors (FGFs) in human DRGs and peripheral nerves, fibroblast growth factor receptors (FGFRs) in mouse fracture callus cells and qRT-PCR of osteogenic genes expression *in vitro*. a. Expression of FGF9 in human sn-RNA seq in relation to expression of markers of nerve type including *SCN10A, NTRK1, NTRK3* and *PVALB*. N=5 human DRGs from N=5 organ donors patients yielding N=2,275 neuronal nuclei^54^. **b** FGF ligands detected in human DRG and sciatic nerves by SOMAscan proteomics assay. Top row: N=16 DRGs and N= 12 sciatic nerve samples from healthy donors. Bottom row: N= 4 DRGs from No Pain condition, N = 9 DRGs from Pain condition. **c** Expression of fibroblast growth factor receptors (FGFRs 1-4) in callus tissue across cell types (left) and across sham versus denervation conditions (right). N=21,920 cells and N=12 mice used for analysis. **d** Osteogenic gene expression (*Runx2*, *Col1a1*, *Sp7*) in periosteal cells was assessed by qRT-PCR. Left: neural CM ± FGFRi experiment. Middle: rmFGF9 treatment experiment. Right: *Fgf9* KD experiment, corresponding to Figure. 4k-n, N=3.

## Main Reference

1 Meltzer, S., Sanago, C., Sharma, N. & Ginty, D. D. The cellular and molecular basis of somatosensory neuron development. Neuron 109, 3736–3757 (2021). hOps://doi.org/10.1016/j.neuron.2021.09.004

2 Noble, A., Qubrosi, R., Cariba, S., Favaro, K. & Payne, S. L. Neural dependency in wound healing and regeneration. Dev Dyn (2023). hOps://doi.org/10.1002/dvdy.650

3 Barrell, K. & Smith, A. G. Peripheral Neuropathy. Med Clin North Am 103, 383–397 (2019). hOps://doi.org/10.1016/j.mcna.2018.10.006

4 Carr, M. J. et al. Mesenchymal Precursor Cells in Adult Nerves Contribute to Mammalian Tissue Repair and Regeneration. Cell Stem Cell 24, 240–256.e249 (2019). hOps://doi.org/10.1016/j.stem.2018.10.024

5 Yin, S. et al. Receptor activity-modifying protein 1 regulates mouse skin fibroblast proliferation via the Gαi3-PKA-CREB-YAP axis. Cell Commun Signal 20, 52 (2022). hOps://doi.org/10.1186/s12964-022-00852-0

6 Huang, S. et al. Lgr6 marks epidermal stem cells with a nerve-dependent role in wound re-epithelialization. Cell Stem Cell 28, 1582–1596.e1586 (2021). hOps://doi.org/10.1016/j.stem.2021.05.007

7 Tarnawski, L. & Olofsson, P. S. Inflammation neuroscience: neuro-immune crosstalk and interfaces. Clin Transl Immunology 10, e1352 (2021). hOps://doi.org/10.1002/c72.1352

8 Tomlinson, R. E. et al. NGF-TrkA Signaling by Sensory Nerves Coordinates the Vascularization and Ossification of Developing Endochondral Bone. Cell Rep 16, 2723–2735 (2016). hOps://doi.org/10.1016/j.celrep.2016.08.002

9 Castañeda-Corral, G. et al. The majority of myelinated and unmyelinated sensory nerve fibers that innervate bone express the tropomyosin receptor kinase A. Neuroscience 178, 196–207 (2011). hOps://doi.org/10.1016/j.neuroscience.2011.01.039

10 Tower, R. J. et al. Spatial transcriptomics reveals a role for sensory nerves in preserving cranial suture patency through modulation of BMP/TGF-β signaling. Proc Natl Acad Sci U S A 118 (2021). hOps://doi.org/10.1073/pnas.2103087118

11 Frampton, J. E. Entrectinib: A Review in NTRK+ Solid Tumours and ROS1+ NSCLC. Drugs 81, 697–708 (2021). hOps://doi.org/10.1007/s40265-021-01503-3

12 Sharma, N. et al. The emergence of transcriptional identity in somatosensory neurons. Nature 577, 392–398 (2020). hOps://doi.org/10.1038/s41586-019-1900-1

13 Wang, K. et al. Single-cell transcriptomic analysis of somatosensory neurons uncovers temporal development of neuropathic pain. Cell Res 31, 904–918 (2021). hOps://doi.org/10.1038/s41422-021-00479-9

14 Li, C., Wang, S., Chen, Y. & Zhang, X. Somatosensory Neuron Typing with High-Coverage Single-Cell RNA Sequencing and Functional Analysis. Neurosci Bull 34, 200–207 (2018). hOps://doi.org/10.1007/s12264-017-0147-9

15 Chan, K. Y. et al. Engineered AAVs for efficient noninvasive gene delivery to the central and peripheral nervous systems. Nat Neurosci 20, 1172–1179 (2017). hOps://doi.org/10.1038/nn.4593

16 Renthal, W. et al. Transcriptional Reprogramming of Distinct Peripheral Sensory Neuron Subtypes aper Axonal Injury. Neuron 108, 128–144.e129 (2020). hOps://doi.org/10.1016/j.neuron.2020.07.026

17 Bachy, I. et al. The transcription factor Cux2 marks development of an A-delta sublineage of TrkA sensory neurons. Dev Biol 360, 77–86 (2011). hOps://doi.org/10.1016/j.ydbio.2011.09.007

18. 18 Chartier, S. R., Mitchell, S. A., Majuta, L. A. & Mantyh, P. W. Immunohistochemical localization of nerve growth factor, tropomyosin receptor kinase A, and p75 in the bone and articular cartilage of the mouse femur. Mol Pain 13, 1744806917745465 (2017). 10.1177/1744806917745465

19 Li, Z. et al. Fracture repair requires TrkA signaling by skeletal sensory nerves. J Clin Invest 129, 5137–5150 (2019). 10.1172/jci128428

20 Li, G., Bunn, J. R., Mushipe, M. T., He, Q. & Chen, X. Effects of pleiotrophin (PTN) over- expression on mouse long bone development, fracture healing and bone repair. Calcif Tissue Int 76, 299–306 (2005). 10.1007/s00223-004-0145-6

21 Li, X. et al. A Single-Cell RNA-Sequencing Analysis of Distinct Subsets of Synovial Macrophages in Rheumatoid Arthritis. DNA Cell Biol 42, 212–222 (2023). 10.1089/dna.2022.0509

22 Himburg, H. A. et al. Pleiotrophin regulates the expansion and regeneration of hematopoietic stem cells. Nat Med 16, 475–482 (2010). 10.1038/nm.2119

23 Clark, D. N., Begg, L. R. & Filiano, A. J. Unique aspects of IFN-γ/STAT1 signaling in neurons. Immunol Rev 311, 187–204 (2022). 10.1111/imr.13092

24 Okamoto, S., Sherman, K., Bai, G. & Lipton, S. A. Effect of the ubiquitous transcription factors, SP1 and MAZ, on NMDA receptor subunit type 1 (NR1) expression during neuronal differentiation. Brain Res Mol Brain Res 107, 89-96 (2002). 10.1016/s0169-328x(02)00440-0

25 Kirjavainen, A. et al. Gata2, Nkx2-2 and Skor2 form a transcription factor network regulating development of a midbrain GABAergic neuron subtype with characteristics of REM-sleep regulatory neurons. Development 149 (2022). 10.1242/dev.200937

26 Gao, X., Daugherty, R. L. & TourtelloOe, W. G. Regulation of low affinity neurotrophin receptor (p75(NTR)) by early growth response (Egr) transcriptional regulators. Mol Cell Neurosci 36, 501–514 (2007). 10.1016/j.mcn.2007.08.013

27 Meyers, K. T. et al. The Immediate Early Gene Egr3 Is Required for Hippocampal Induction of Bdnf by Electroconvulsive Stimulation. Front Behav Neurosci 12, 92 (2018). 10.3389/fnbeh.2018.00092

28 Wu, F. & Li, C. KLF2 up-regulates IRF4/HDAC7 to protect neonatal rats from hypoxic- ischemic brain damage. Cell Death Discov 8, 41 (2022). 10.1038/s41420-022-00813-z

29 Lei, Z., Stone, S. & Lin, W. Detection of PERK Signaling in the Central Nervous System. Methods Mol Biol 2378, 233–245 (2022). 10.1007/978-1-0716-1732-8_15

30 Chen, X. et al. A chemical-genetic approach to studying neurotrophin signaling. Neuron 46, 13–21 (2005). 10.1016/j.neuron.2005.03.009

31 Huang, S. et al. Lymph nodes are innervated by a unique population of sensory neurons with immunomodulatory potential. Cell 184, 441–459. e425 (2021).

32 Jimenez-Andrade, J. M. et al. A phenotypically restricted set of primary afferent nerve fibers innervate the bone versus skin: therapeutic opportunity for treating skeletal pain. Bone 46, 306–313 (2010). 10.1016/j.bone.2009.09.013

33. 33 Oostinga, D., Steverink, J. G., van Wijck, A. J. M. & Verlaan, J. J. An understanding of bone pain: A narrative review. Bone 134, 115272 (2020). 10.1016/j.bone.2020.115272

34 Liu, S. et al. A neuroanatomical basis for electroacupuncture to drive the vagal–adrenal axis. Nature 598, 641–645 (2021).

35 Wang, K., Cai, B., Song, Y., Chen, Y. & Zhang, X. Somatosensory neuron types and their neural networks as revealed via single-cell transcriptomics. Trends Neurosci 46, 654–666 (2023). 10.1016/j.7ns.2023.05.005

36 Handler, A. et al. Three-dimensional reconstructions of mechanosensory end organs suggest a unifying mechanism underlying dynamic, light touch. Neuron 111, 3211–3229. e3219 (2023).

37 Zampieri, N. & de Nooij, J. C. Regulating muscle spindle and Golgi tendon organ proprioceptor phenotypes. Curr Opin Physiol 19, 204–210 (2021). 10.1016/j.cophys.2020.11.001

38 Blecher, R. et al. The Proprioceptive System Regulates Morphologic Restoration of Fractured Bones. Cell Rep 20, 1775–1783 (2017). 10.1016/j.celrep.2017.07.073

39 Gong, L. et al. Global analysis of transcriptome in dorsal root ganglia following peripheral nerve injury in rats. Biochem Biophys Res Commun 478, 206–212 (2016). 10.1016/j.bbrc.2016.07.067

40 Korczeniewska, O. A. et al. Time-Course Progression of Whole Transcriptome Expression Changes of Trigeminal Ganglia Compared to Dorsal Root Ganglia in Rats Exposed to Nerve Injury. J Pain 25, 101–117 (2024). 10.1016/j.jpain.2023.07.024

41 Todd, T. J. On the process of reproduction of the members of the aquatic salamander. *Quarterly Journal of Science*, Literatures and the Arts 16, 84–86 (1823).

42 Farkas, J. E. & Monaghan, J. R. A brief history of the study of nerve dependent regeneration. Neurogenesis (AusIn) 4, e1302216 (2017). 10.1080/23262133.2017.1302216

43 Mahmoud, A. I. et al. Nerves Regulate Cardiomyocyte Proliferation and Heart Regeneration. Dev Cell 34, 387–399 (2015). 10.1016/j.devcel.2015.06.017

44 Johnston, A. P. et al. Dedifferentiated Schwann Cell Precursors Secreting Paracrine Factors Are Required for Regeneration of the Mammalian Digit Tip. Cell Stem Cell 19, 433–448 (2016). 10.1016/j.stem.2016.06.002

45 Roosterman, D., Goerge, T., Schneider, S. W., BunneO, N. W. & Steinhoff, M. Neuronal control of skin function: the skin as a neuroimmunoendocrine organ. Physiol Rev 86, 1309–1379 (2006). 10.1152/physrev.00026.2005

46 Pei, F. et al. Sensory nerve niche regulates mesenchymal stem cell homeostasis via FGF/mTOR/autophagy axis. Nat Commun 14, 344 (2023). 10.1038/s41467-023-35977-4

47 Zhao, H. et al. Secretion of shh by a neurovascular bundle niche supports mesenchymal stem cell homeostasis in the adult mouse incisor. Cell Stem Cell 14, 160–173 (2014). 10.1016/j.stem.2013.12.013

48 Qin, Q. et al. Neuron-to-vessel signaling is a required feature of aberrant stem cell commitment aper sop 7ssue trauma. Bone Res 10, 43 (2022). 10.1038/s41413-022-00216-x

49 Chu, C., Artis, D. & Chiu, I. M. Neuro-immune interactions in the tissues. Immunity 52, 464-474 (2020).

50 Qin, Q. et al. Neurovascular coupling in bone regeneration. Exp Mol Med 54, 1844–1849 (2022). 10.1038/s12276-022-00899-6

51 Wentz, L., Liu, P. Y., Haymes, E. & Ilich, J. Z. Females have a greater incidence of stress fractures than males in both military and athletic populations: a systemic review. Mil Med 176, 420–430 (2011). 10.7205/milmed-d-10-00322

52 Rollman, G. B. & Lautenbacher, S. Sex differences in musculoskeletal pain. Clin J Pain 17, 20–24 (2001). 10.1097/00002508-200103000-00004

53 Tavares-Ferreira, D. et al. Spatial transcriptomics of dorsal root ganglia identifies molecular signatures of human nociceptors. Sci Transl Med 14, eabj8186 (2022). 10.1126/scitranslmed.abj8186

54 Bhuiyan, S. A. et al. Harmonized cross-species cell atlases of trigeminal and dorsal root ganglia. bioRxiv (2023). 10.1101/2023.07.04.547740

## Methods References

1 Li, Z. et al. Fracture repair requires TrkA signaling by skeletal sensory nerves. J Clin Invest 129, 5137–5150 (2019). 10.1172/jci128428

2 Tower, R. J. et al. Spatial transcriptomics reveals a role for sensory nerves in preserving cranial suture patency through modulation of BMP/TGF-β signaling. Proc Natl Acad Sci U S A 118 (2021). 10.1073/pnas.2103087118

3 Chen, X. et al. A chemical-genetic approach to studying neurotrophin signaling. Neuron 46, 13–21 (2005). 10.1016/j.neuron.2005.03.009

4 Chan, K. Y. et al. Engineered AAVs for efficient noninvasive gene delivery to the central and peripheral nervous systems. Nat Neurosci 20, 1172–1179 (2017). 10.1038/nn.4593

5 Renier, N. et al. iDISCO: a simple, rapid method to immunolabel large tissue samples for volume imaging. Cell 159, 896–910 (2014). 10.1016/j.cell.2014.10.010

6 Martinez, M. D., Schmid, G. J., McKenzie, J. A., Ornitz, D. M. & Silva, M. J. Healing of non-displaced fractures produced by fatigue loading of the mouse ulna. Bone 46, 1604–1612 (2010). 10.1016/j.bone.2010.02.030

7 Bouxsein, M. L. et al. Guidelines for assessment of bone microstructure in rodents using micro-computed tomography. J Bone Miner Res 25, 1468–1486 (2010). 10.1002/jbmr.141

8 Zeisel, A. et al. Molecular Architecture of the Mouse Nervous System. Cell 174, 999–1014.e1022 (2018). 10.1016/j.cell.2018.06.021

9 Hao, Y. et al. Integrated analysis of multimodal single-cell data. Cell 184, 3573–3587.e3529 (2021). 10.1016/j.cell.2021.04.048

10 McGinnis, C. S., Murrow, L. M. & Gartner, Z. J. DoubletFinder: Doublet Detection in Single-Cell RNA Sequencing Data Using Artificial Nearest Neighbors. Cell Syst 8, 329–337.e324 (2019). 10.1016/j.cels.2019.03.003

11 Korsunsky, I. et al. Fast, sensitive and accurate integration of single-cell data with Harmony. Nat Methods 16, 1289–1296 (2019). 10.1038/s41592-019-0619-0

12 Reimand, J. et al. Pathway enrichment analysis and visualization of omics data using g:Profiler, GSEA, Cytoscape and EnrichmentMap. Nat Protoc 14, 482–517 (2019). 10.1038/s41596-018-0103-9

13 Aibar, S. et al. SCENIC: single-cell regulatory network inference and clustering. Nat Methods 14, 1083–1086 (2017). 10.1038/nmeth.4463

14 Wu, T. et al. clusterProfiler 4.0: A universal enrichment tool for interpreting omics data. Innovation (Camb) 2, 100141 (2021). 10.1016/j.xinn.2021.100141

15 Jin, S. et al. Inference and analysis of cell-cell communication using CellChat. Nat Commun 12, 1088 (2021). 10.1038/s41467-021-21246-9

16 Browaeys, R., Saelens, W. & Saeys, Y. NicheNet: modeling intercellular communication by linking ligands to target genes. Nat Methods 17, 159–162 (2020). 10.1038/s41592-019-0667-5

17 Bhuiyan, S. A. et al. Harmonized cross-species cell atlases of trigeminal and dorsal root ganglia. bioRxiv (2023). 10.1101/2023.07.04.547740

18 Schindelin, J., et al. Fiji: an open-source platform for biological-image analysis. Nat Methods 9, 676-682 (2012). 10.1038/nmeth.2019

19 Xu, J. et al. PDGFRα reporter activity identifies periosteal progenitor cells critical for bone formation and fracture repair. Bone Res 10, 7 (2022). 10.1038/s41413-021-00176-8

20 Wang, X. W. et al. Lin28 Signaling Supports Mammalian PNS and CNS Axon Regeneration. Cell Rep 24, 2540–2552.e2546 (2018). 10.1016/j.celrep.2018.07.105

21 Thottappillil, N. et al. ZIC1 dictates osteogenesis versus adipogenesis in human mesenchymal progenitor cells via a Hedgehog dependent mechanism. Stem Cells (2023). 10.1093/stmcls/sxad047

22 Kim, C. H. et al. Stability and reproducibility of proteomic profiles measured with an aptamer-based platform. Sci Rep 8, 8382 (2018). 10.1038/s41598-018-26640-w

